# SRARec: A program for detecting recombination in sequencing reads and its application to uncover recombination patterns in SARS-CoV-2 and HIV-1

**DOI:** 10.64898/2026.07.30.741747

**Authors:** Luis D. González-Vázquez, Paula Iglesias-Rivas, Miguel Arenas, Darren P. Martin

## Abstract

The detection of recombination using consensus genome sequences has key limitations including failure to consider rare genetic variants and misidentification of artifactually assembled genome chimaeras as biological recombinants. However, commonly used recombination detection tools are not designed to directly analyse sequencing read data. Here, we present *SRARec*, a recombination detection tool that operates directly on raw reads. *SRARec* identifies polymorphic sites and applies the four-gamete test to detect recombination at the read level. The software incorporates mapping and quality filters and can analyse large repositories of raw sequencing data. Simulation validations showed that, given sufficient sequence diversity, *SRARec* can accurately detect recombination breakpoints. The consideration of rare variants makes *SRARec* particularly useful for detecting recombination in intra-host viral populations. Therefore, we applied the tool to 601,045 SARS-CoV-2 and 4,999 HIV-1 read datasets from the Sequence Read Archive (SRA, NCBI), enabling unprecedented genome-wide screening of intra-host recombination breakpoint signals at read-level resolution. Aggregating across all analysed datasets, the distribution of detected recombination breakpoint counts along genomes differed between these viruses, with pervasive breakpoint signals detectable in HIV-1 and sporadic clustered breakpoint hotspots in SARS-CoV-2.

**Graphical abstract:** 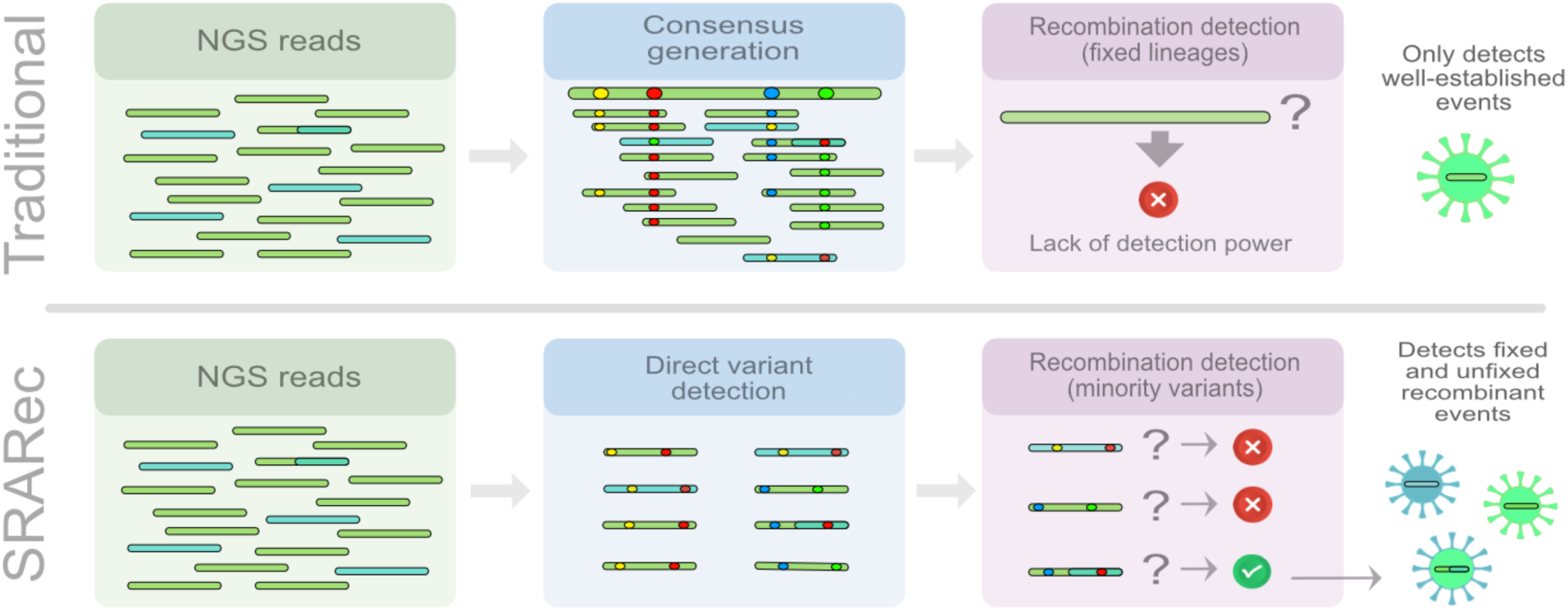

## Introduction

Genetic recombination in viruses can generate genetic variants with altered transmissibility, pathogenicity, immune escape and drug resistance [1–4], potentially impacting decisions made by clinicians and public health officials [5]. Further, if unaccounted for, recombination can mislead certain of the evolutionary and molecular epidemiological inferences that can be drawn from nucleotide sequence datasets [6–10].

Recombination has therefore been broadly studied and numerous methods (i.e., [11–22]) have been developed for detecting recombination in genetic data. However, the vast majority of these methods have been designed to analyse consensus sequences that each represent the “average” of a viral population within the DNA or RNA sample from which the sequence was determined. Because these consensus sequences squash intra-host diversity into a single majority-rule genome, they can obscure real evolutionary events that might otherwise have been detectable if low-frequency variants had also been considered. However, existing recombination analysis software cannot directly process raw high-throughput sequencing (HTS) reads, leaving a major biological and technical blind spot.

Further, with short read sequences determined from samples containing mixtures of genetically distinct genomes, there is no reliable way to assemble the individual reads into genome-length contigs without the risk of generating false, artifactual recombinants that contain sets of polymorphisms that never actually co-existed in a single genome (Figure 1) [23–26]. This limitation is further compounded by experimental artifacts such as RT-PCR generated chimeras and amplicon biases, which can either mimic or obscure genuine biological recombination events (Figure 1) [27,28].

**Figure 1.**
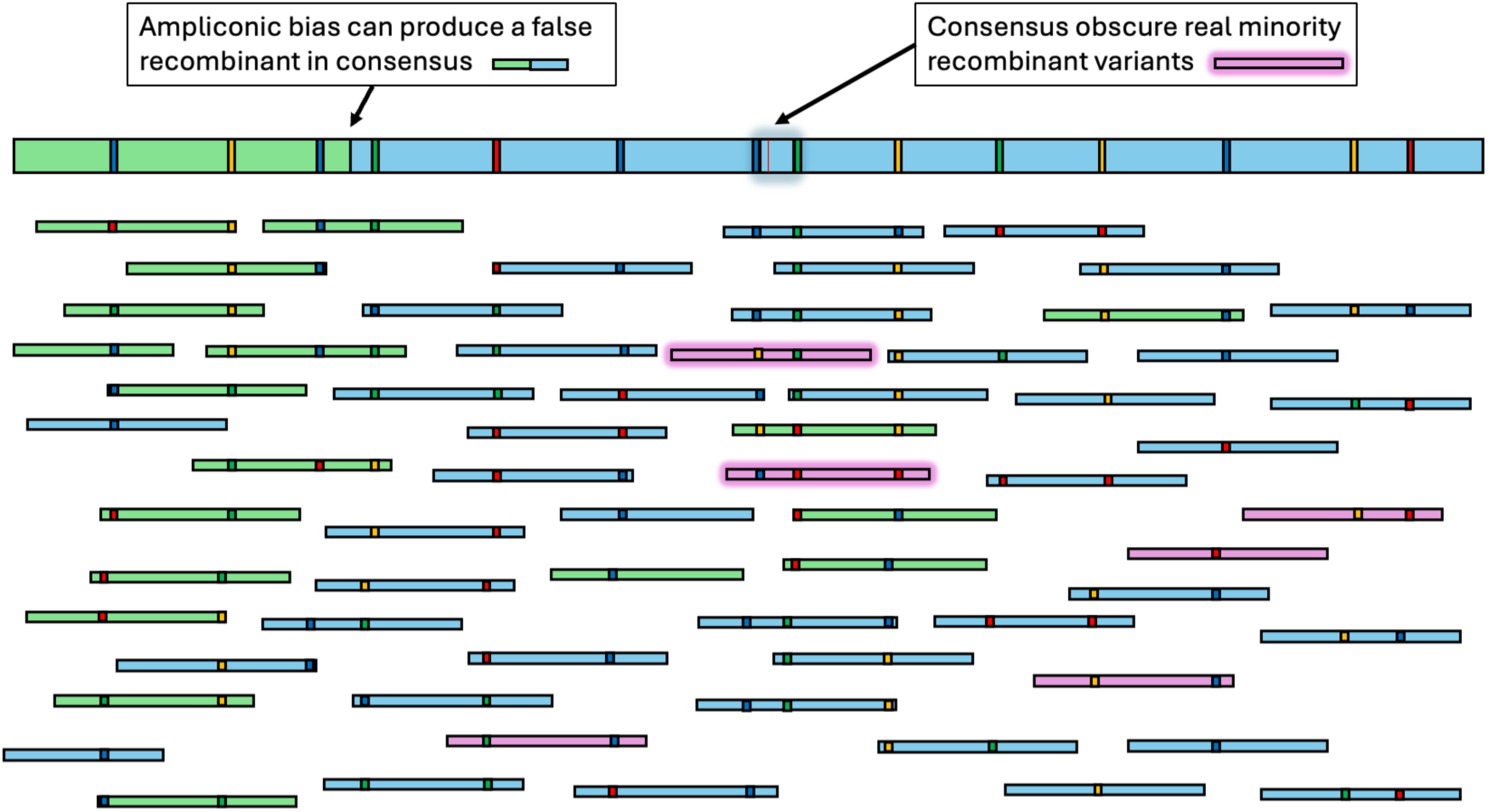
How consensus genomes can create false recombinants and mask true intra-host recombination. Reads from multiple co-circulating variants (green, blue and purple) are aligned. In the first quarter of the alignment amplicons from the green variant predominate, whereas in the remaining three quarters of the genome amplicons from the blue variant predominate. A consensus sequence built by selecting the majority allele at each site therefore becomes a mosaic of the dominant genotypes across regions, yielding an artificial recombinant-like sequence that can be detected as recombinant by consensus-based recombination detection methods even though no read-level haplotypes support such a breakpoint. Conversely, a genuine homologous recombination event between variants (illustrated by the emergence of the purple variant, carrying a combination of mutations from the blue/green and yellow/red backgrounds) may remain undetected in consensus analyses when the recombinant is at low abundance. In particular, the diagnostic haplotype combination is diluted by the more prevalent variants and is not represented in the consensus, thereby obscuring true intra-host recombination.

To sidestep assembly-driven artifacts, short reads can be treated as short haplotypes to directly evaluate linkage. We have noted that, from a population genetics perspective, observing all four haplotypes for pairs of biallelic sites (the four-gamete test, FGT) [29] linked within a single sequencing read can inform recombination detection without requiring full-length genome assemblies. While adaptations of the FGT have been implemented in tools like *PHITest* [14], *LDHat* [13,15] and a variety of other linkage disequilibrium-based recombination-rate estimation methods [30–33], these applications still rely on consensus sequence inputs and remain vulnerable to the same whole genome sequence assembly errors: for example, hybrid severe acute respiratory syndrome coronavirus 2 (SARS-CoV-2) consensus genomes that appeared to be Delta-Omicron recombinants due to the primer set used for sequencing [27]. Raw sequencing reads can, however, include invaluable genetic signatures of recombination trapped in low-frequency variants, which is key for appropriately analysing recombination occurring within intra-host viral populations [34]. To demonstrate the utility of read-level FGT screening across divergent viral dynamics, here we examine two contrasting cases of general interest: SARS-CoV-2 and Human immunodeficiency virus 1 (HIV-1).

Recombination occurring among the coronavirus progenitors of SARS-CoV-2 within bats and/or other mammalian species was likely involved in the origin of SARS-CoV-2 [35,36]. Mechanistically, recombination in coronaviruses is generally understood to occur via a copy-choice mechanism mediated by RNA-dependent RNA polymerase (RdRp) template switching during RNA synthesis (Figure 2A) [37–39]. In SARS-CoV-2, this template-switching capacity is also central to discontinuous transcription, whereby RdRp switches templates during the generation of subgenomic RNAs, guided by transcription regulatory sequences (TRS) [40]. Although this provides a plausible route by which TRS-associated regions could facilitate recombination between co-occurring viral genomes, the extent to which TRS sites act as preferential recombination breakpoints should be considered a possible mechanism rather than a fully established general rule [2,41].

**Figure 2.**
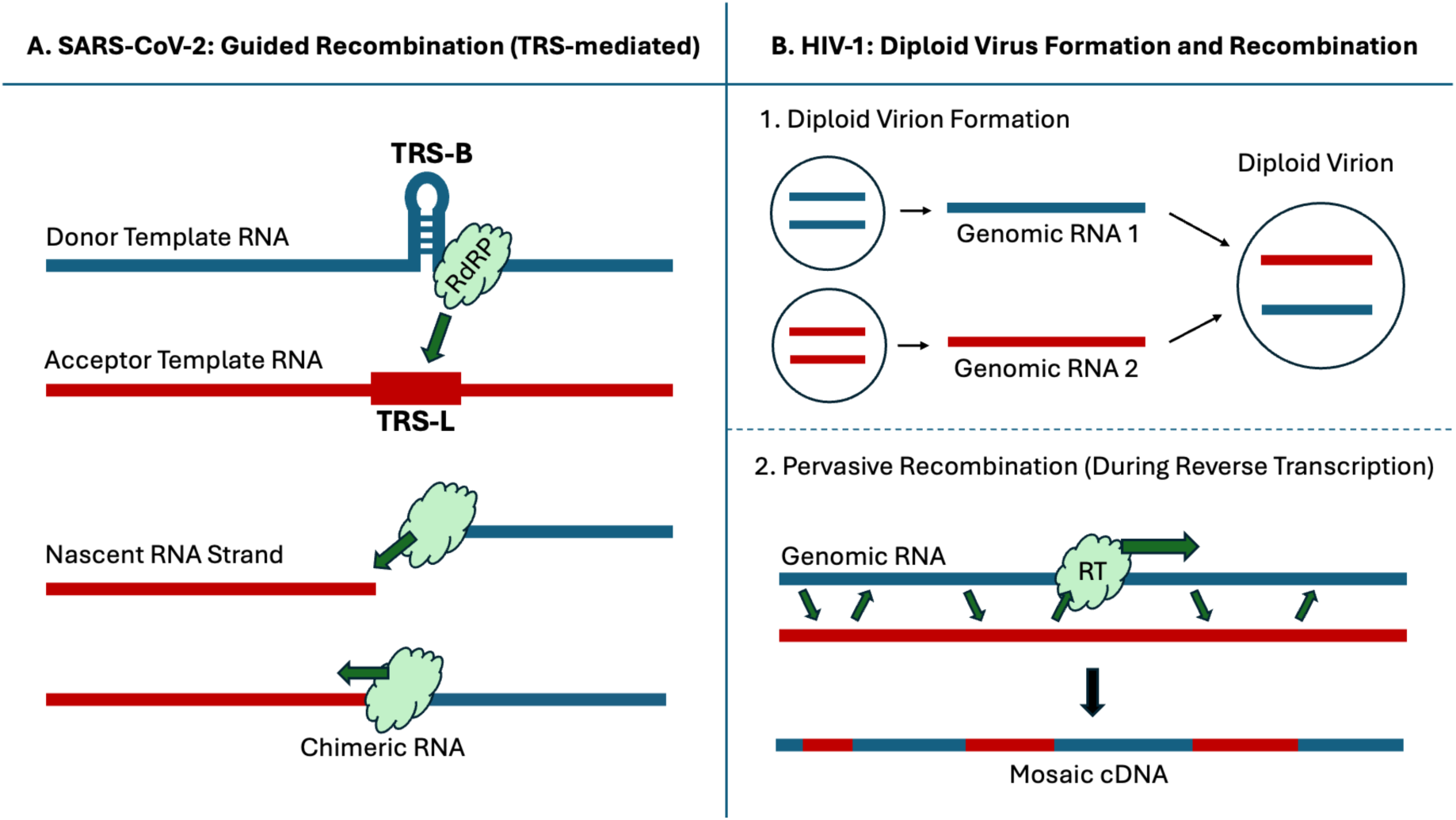
Contrasting recombination mechanisms in SARS-CoV-2 and HIV-1. This schematic summarizes two routes by which RNA viruses can generate chimeric genomes through polymerase template switching. (A) SARS-CoV-2. Coronavirus recombination is generally attributed to RNA-dependent RNA polymerase (RdRp) template switching during RNA synthesis. In SARS-CoV-2, template switching also occurs during discontinuous transcription, when the RdRp switches from a body transcription-regulatory sequence (TRS-B) to the leader TRS (TRS-L), generating subgenomic RNAs. Under coinfection, analogous template-switching events between distinct genomic RNA templates could contribute to the generation of recombinant genomes. However, the role of TRS-associated regions as preferential recombination breakpoints should be interpreted as a plausible or proposed mechanism rather than as a universally demonstrated rule. Recombination can also occur during genome replication at non-TRS sites. (B) HIV-1. HIV-1 recombination is facilitated by the packaging of two RNA genome copies within each virion. Coinfection by different viral lineages can generate heterozygous virions containing two distinct parental RNA genomes. During reverse transcription, reverse transcriptase can switch repeatedly between these co-packaged templates, producing recombinant proviral DNA.

Early in the pandemic, reliably detecting recombination at the global population-scale was almost impossible because SARS-CoV-2 diversity was too low [42], especially in comparison to other RNA viruses such as HIV-1 [43]. As SARS-CoV-2 genetic diversity increased over time, recombination between genetically divergent lineages became more easily detectable at the population-scale, with inter-lineage recombinants now being demarcated with an “X” lineage designation prefix [44–53]. At the intra-host scale, however, SARS-CoV-2 recombination remains difficult to detect because narrow transmission bottlenecks and short infection durations mean that even deep sequencing commonly fails to reveal sufficient intra-host genetic diversity for recombination to be detectable [54–56]. While circulating recombinant lineages are detected and tracked using consensus genome sequences, these recombinants must have ultimately originated from individual recombination events occurring within individual coinfected cells. To date, only a few studies have reported direct, read-level evidence of SARS-CoV-2 intra-host recombination in specific cases of coinfection or superinfection [47,51,57,58]. Because global surveillance efforts have largely relied on the analysis of these low-diversity consensus sequences, we hypothesized that the recombinants characterized so far represent only a small fraction of the true recombination signals present in the voluminous publicly available SARS-CoV-2 raw read-level data that has accumulated since the start of the pandemic [34,53,59].

Relative to within-host SARS-CoV-2 populations, within-host HIV-1 populations usually show substantially higher degrees of genetic diversity, with intra-host recombination being readily detectable [3]. Mechanistically, HIV-1 recombination is enabled by the encapsidation of two RNA genome copies within each virion, such that coinfection can generate heterozygous virions containing two different parental RNAs (Figure 2B) [60,61]. During reverse transcription, HIV-1 reverse transcriptase frequently switches between these co-packaged templates, producing recombinant DNA genomes as a routine outcome of replication [62–64]. As a consequence, recombination in HIV-1 can occur at rates comparable to mutation, and may even exceed mutation rates in some contexts, making it a major driver of viral diversification [3,62]. Consistent with its major contribution to HIV-1 evolution, approximately 16.7% of HIV-1 infections globally are attributable to inter-subtype recombinants referred to as circulating recombinant forms (CRFs) [2,65,66]. A particular concern is that recombination can combine intra-host antiretroviral drug-resistance mutations from different lineages into single multi-drug-resistant genomes, accelerating viral adaptation under therapy [3,62]. However, as with SARS-CoV-2, the use of single consensus genome sequences from sampled infections provides little information about the ongoing recombination processes within individual patients. In this regard, analysing raw sequence read data from individual HIV-1 infections could likely reveal recombination breakpoints in multiple low-frequency recombinants within individual infections and, when aggregated across multiple datasets from different patients, reveal site-to-site variation in recombination frequencies.

In order to detect a genuine biological recombinant genome [3], the genome must have been viable enough that its progeny expanded to a frequency high enough to be sampled, and the relevant parental and recombinant haplotypes must have been present at the time points and in the anatomical compartments that were sequenced [2,28,56]. As a result, recombinant lineages that are circulating at the population-level and that are eventually sampled represent a highly filtered subset of all recombination events, shaped not only by selection for viability and replication but also by the randomness of transmission and sampling. Intra-host studies can partially bypass these selection and transmission-driven filters by revealing recombinant haplotypes before they are selected (or purged) by onward spread, even though detection still depends on within-host selection processes and sampling depth [55,67,68].

As previously noted and given the potential evolutionary impacts of recombination within individual infections, detecting and determining the patterns of intra-host recombination is a relevant endeavor. Besides revealing real-time interactions among variants and identifying the existing variant landscape within a population, when studied across short-read data from thousands of individual infections, these analyses could potentially reveal the innate mechanistic predispositions for recombination to occur at certain genome sites. They could also yield actionable information on the statistical probabilities of recombination assembling multiple drug resistance or immune-evasion mutations within single genomes.

Here we present SRARec, software to detect recombination directly from raw reads by identifying polymorphic site pairs and applying read-level FGTs under stringent mapping and quality filters. By accounting for the co-occurrence of minority variants within individual short reads, SRARec both avoids being confounded by artifactual assembly-generated recombination signals and recovers weak recombination signals that would otherwise be invisible to, and which could also potentially mislead, recombination detection based on assembled consensus sequences. Running on either the command line (CLI) or via a graphical user interface (GUI), SRARec can infer recombination breakpoint distributions across massive repositories of read sequencing data collected from hundreds of thousands of individual viral infections using a variety of sequencing platforms. As a demonstration, we analysed 601,045 SARS-CoV-2 and 4,999 HIV-1 single read archive (SRA) accessions to generate genome-wide maps illustrating site-to-site variations in intra-host recombination frequencies. Interestingly, SRARec is compatible with the sorts of high throughput sequence data processing pipelines that are routinely used during viral genomic surveillance studies, enabling the automated flagging of likely recombinant reads for further analysis or verification, and thus accelerating the early detection of recombinationally assembled viral variants that can be of potential epidemiological relevance.

## Methods

### The SRARec method and its software implementation

SRARec is computer software developed in Python 3 and distributed as open source. Its source code, along with the GUI, detailed documentation, and ready-to-use examples, is freely available from the GitHub repository at https://github.com/ldgonzalezvazquez/SRARec/.

SRARec allows the analysis of sequencing reads generated by the three major sequencing platforms, including Illumina, Oxford Nanopore Technologies (ONT), and Pacific Biosciences (PacBio), with both single-end (SE) and paired-end (PE) data. It can also directly obtain data from the NCBI Sequence Read Archive (NCBI–SRA) database [69,70]. Thus, SRARec allows users to choose between *(1)* a local mode by specifying one FASTQ file (for SE data), two FASTQ files (for PE data) or a BAM file, and *(2)* a NCBI– SRA mode by entering a database accession code (standard format SRR/ERR/DRR).

SRARec implements two pipeline schemes: (*i*) a recommended ‘slow’ mode for low-diversity or low-coverage datasets (trimming, mapping, polymorphism calling and recombination detection), and (*ii*) a ‘fast’ mode for high-diversity and high-coverage datasets (mapping, polymorphism calling, read subset trimming and remapping, polymorphism calling and recombination detection). Before trimming, orphan reads are removed from paired-end runs with the *BBMap* module, *Repair.sh* [71]. Trimming is carried out with *cutadapt* [72]. The default configuration includes an adapter database from *BBMap*, but users may specify a custom adapter file when adapter sequences are known.

The reads are then mapped using *BWA-MEM* [73,74] for Illumina and *minimap2* [75,76] for ONT and PacBio. Duplicates are then removed and several filters are applied to the mapping (MAPQ > 30; exclusion of unmapped reads; base quality above 20; elimination of reads that map below 90% of their length or map in more than one place), in addition to ordering and indexing [77,78]. From the resulting BAM file, mapping information is extracted with *SAMtools mpileup* [77–79] and then the alignment is processed only in the positions that have two or more alleles and N or more supporting reads (by default, N=10) for the minor allele.

For highly divergent datasets in which direct mapping to a single reference can be suboptimal (e.g., HIV-1), SRARec can be run using a hybrid “*de novo* + reference-guided” preprocessing strategy to generate a sample-specific consensus while retaining reference-based coordinates. Briefly, the sequencing reads are first assembled *de novo* into contigs using *SPAdes* [80], and the resulting contigs are then used within a reference-guided refinement workflow implemented in *Shiver* [81] to obtain a consensus sequence and corresponding alignments anchored to the chosen reference. The downstream SRARec analysis is then performed on the resulting reference-anchored BAM file using the filtering and procedures previously described.

All sequencing reads overlapping at least two polymorphic sites in the BAM files, which resulted from the previously indicated methodologies, are then extracted. Each read is examined to determine its allele state at every overlapping polymorphic position (Figure 3), including insertions and deletions (indels), which are recorded at their leftmost (left-aligned) coordinate.

**Figure 3.**
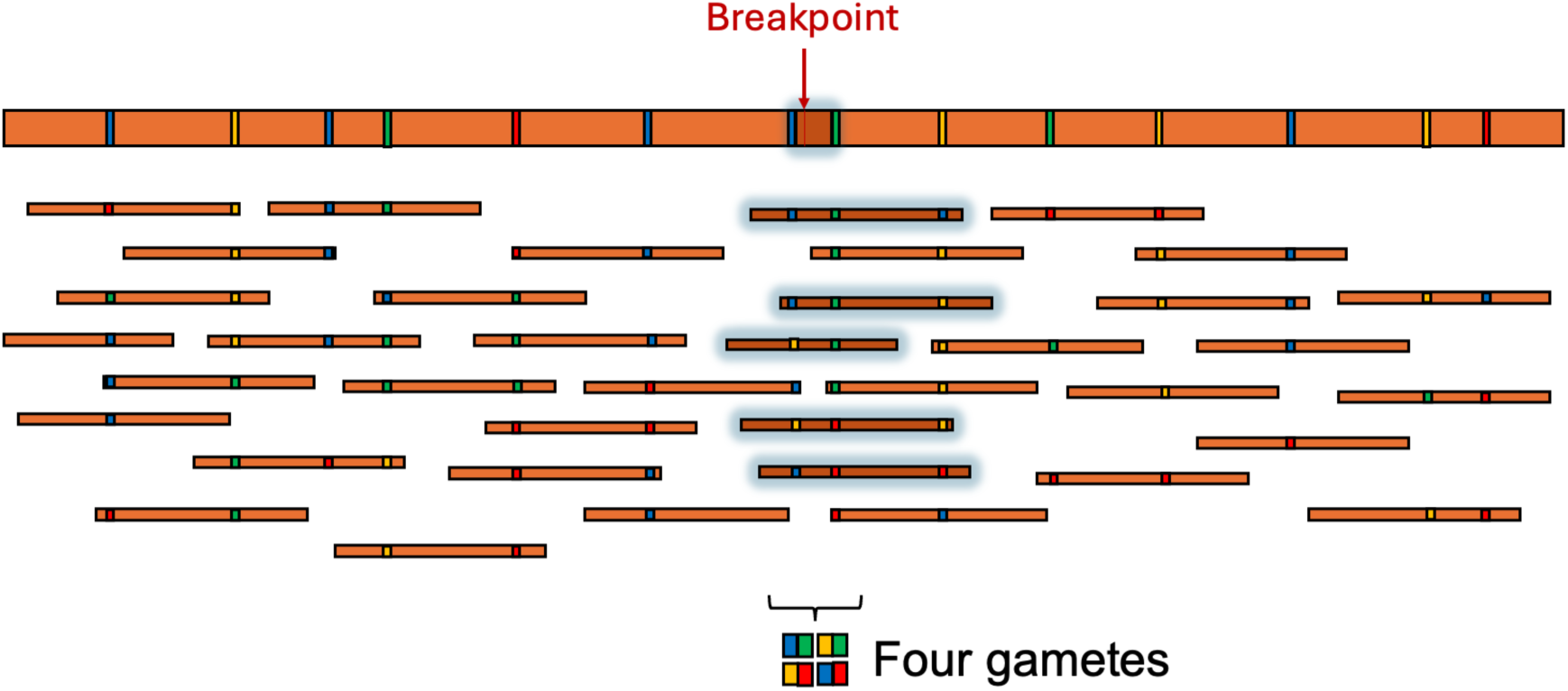
Read-level haplotypes used for four-gamete test screening. Schematic illustration of how SRARec explores linkage between polymorphic sites using individual reads. From the alignment, SRARec extracts only reads that overlap at least two polymorphic positions, then records the allele state carried by each read at all overlapping sites to define read-level haplotypes. Haplotype combinations observed across reads are subsequently evaluated for signals consistent with recombination using the four-gamete test.

Next, the set of detected polymorphisms is used to generate analysis windows defined with the maximum read size mapped from each polymorphic position (Figure 4). Indeed, within each window, analysis groups are defined based on the allele-frequency profiles observed at each polymorphic position. In particular, positions are assigned to the same group when at least one allele-frequency component differs by <5% from the corresponding group mean (Figure 5), and a position is allowed to belong to multiple groups. For each group within each window, comparisons are made starting with the pair of positions that are farthest apart. If a comparison is negative, the remaining comparisons in that set are considered as negative. If it is positive, the breakpoint location is refined by comparing the two most distant polymorphic positions with the median position of the group. This procedure is then applied recursively to all positive pairwise comparisons, allowing the breakpoint position to be progressively narrowed down (Figures 4 and 5). In the case of an even number of positions, the left position is compared with the position to the right of the median and *vice versa*, in addition to comparing both positions on either side of the median with each other. To account for recombination breakpoints located between groups within the same window, adjacent positions assigned to different groups are also compared.

**Figure 4.**
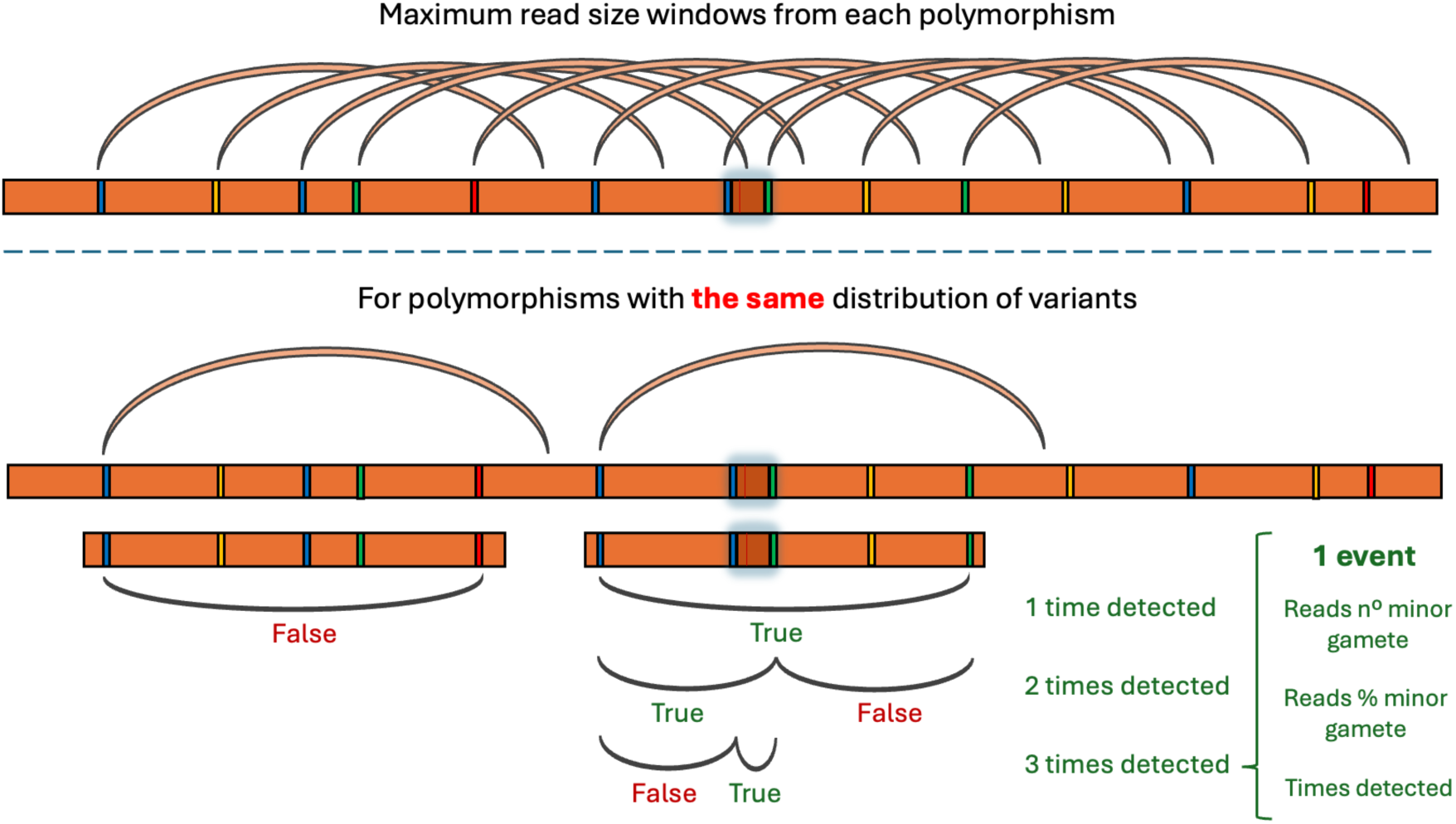
Window construction and coarse-to-fine search within windows. Schematic definition of analysis windows in SRARec and iteratively refinement of candidate signals. For each polymorphic position, a forward window is created spanning the maximum mapped read length from that position, thereby restricting comparisons to site pairs that can be linked by individual reads. Within each window, SRARec initiates testing with the two polymorphic sites that are farthest apart (window extremes). When a positive signal is detected (i.e. all four possible gametes at the site pair are found), the search is refined by testing progressively more central sites (recursive “divide-and-refine” from the extremes toward the median), whereas a negative result at the extremes terminates the search for that set of comparisons. The figure also illustrates that multiple overlapping windows can contribute to repeated detections of the same underlying event, which are recorded for downstream summarization.

**Figure 5.**
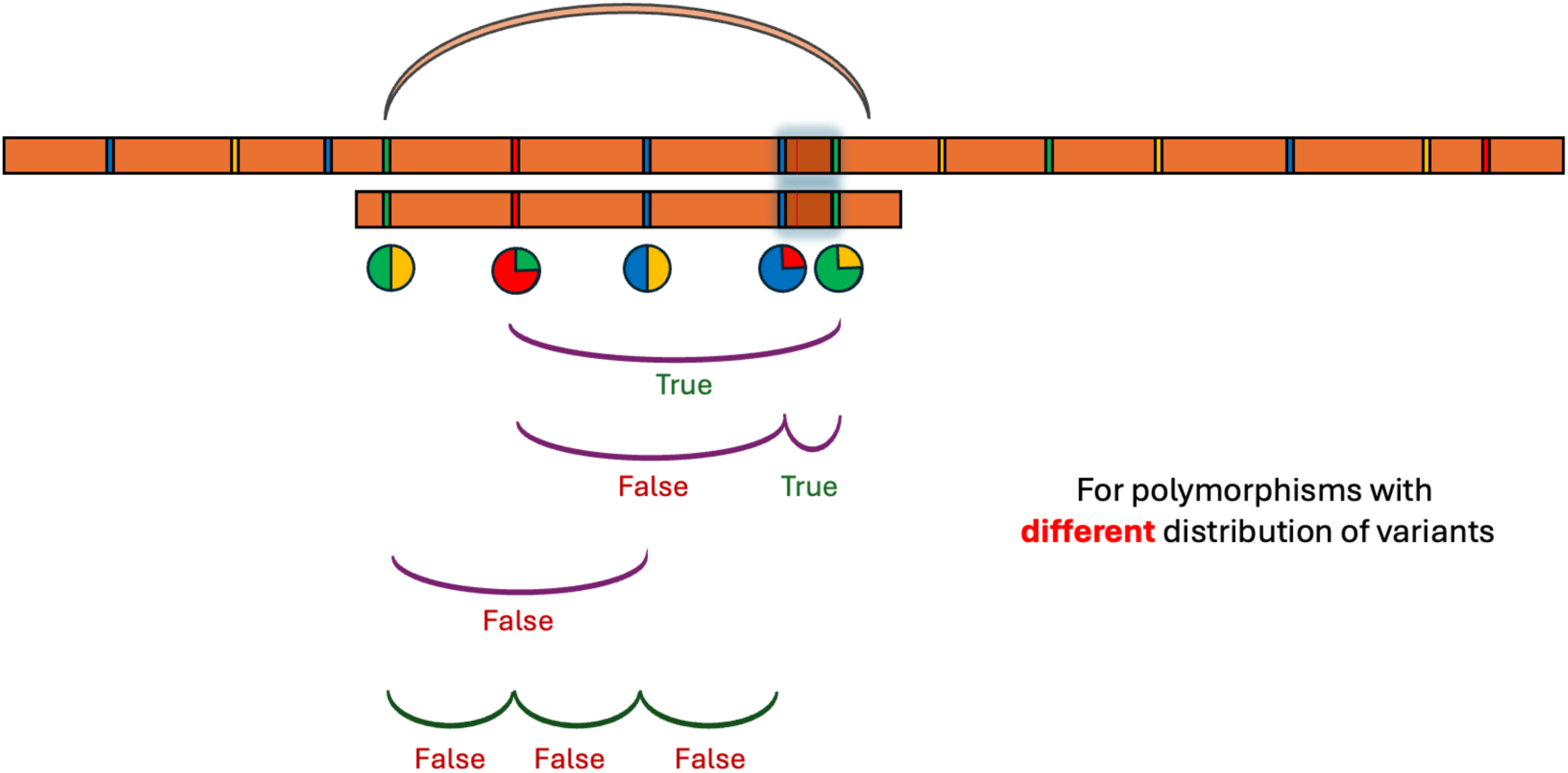
Frequency-based grouping of polymorphic sites within windows. Schematic of how SRARec forms analysis groups from allele-frequency profiles observed at each polymorphic position within a window. Positions that have similar allele-frequency vectors (within a fixed tolerance) are assigned to the same group and analysed together using the refinement strategy shown in Figure 4. Because similarity is assessed relative to group profiles, a given position may belong to more than one group. In addition to within-group testing, SRARec performs comparisons between groups by evaluating all adjacent pairs of polymorphic positions that do not share a group. Results from both within-group refinement and between-group adjacent-position comparisons are recorded and annotated for downstream summarization and further refinement.

For each pairwise site comparison, SRARec evaluates whether the four-gamete test is satisfied based on the haplotype combinations observed across individual reads (Figure 3). In the absence of filtering, the test is positive when reads overlapping the same two polymorphic sites (A and B) support all four haplotypes: A=X/B=X, A=X/B=Y, A=Y/B=X, and A=Y/B=Y (all four “gametes”), a pattern that is incompatible with a simple non-recombining scenario. By default, filters are applied to guarantee events that have at least five reads per gamete and 5% frequency among all possible gametes. These filters can be changed during execution, but we recommend at least these levels to minimize the risk of false positives due to PCR chimeras, according to [51].

The main output of SRARec is a table of site pairs A and B that satisfy the four-gamete criterion and are therefore flagged as pair records, each representing a recombination signal. For each flagged A and B pair, the table reports the alleles defining the four haplotypes, the observed frequency of each gamete, the number of supporting reads, and how often the same event is recovered when evaluated within larger comparison sets. The same output also records A and B pairs that do not meet the criterion. In addition, SRARec outputs the start and end coordinates of the reads that contributed to both results, enabling downstream assessment of potential amplicon-related biases.

To summarize these discrete pairwise A-B calls as a continuous genome-wide profile (Figure 6), the tool accumulates two per-position tracks, T(x) and F(x), representing the total “True” and “False” support at nucleotide resolution. Each A-B pair record is converted into a per-nucleotide site contribution by spreading a unit weight uniformly across the genomic interval between site positions A and B. Specifically, a record spanning the interval between nucleotide sites A and B contributes w=1/(B–A) to every nucleotide position between sites A and B. Contributions from “True” records are accumulated into T(x) and those from “False” records into F(x). When multiple records overlap the same position within a given individual sample, the method retains the most informative signal by prioritizing “True” over “False” and, among “True” records, the highest contribution is kept for that position. This normalization ensures that distant A-B pairs do not dominate solely by covering more bases. A positional four-gamete test ratio (FGT ratio) is then computed as T(x)/(T(x)+F(x)).

**Figure 6.**
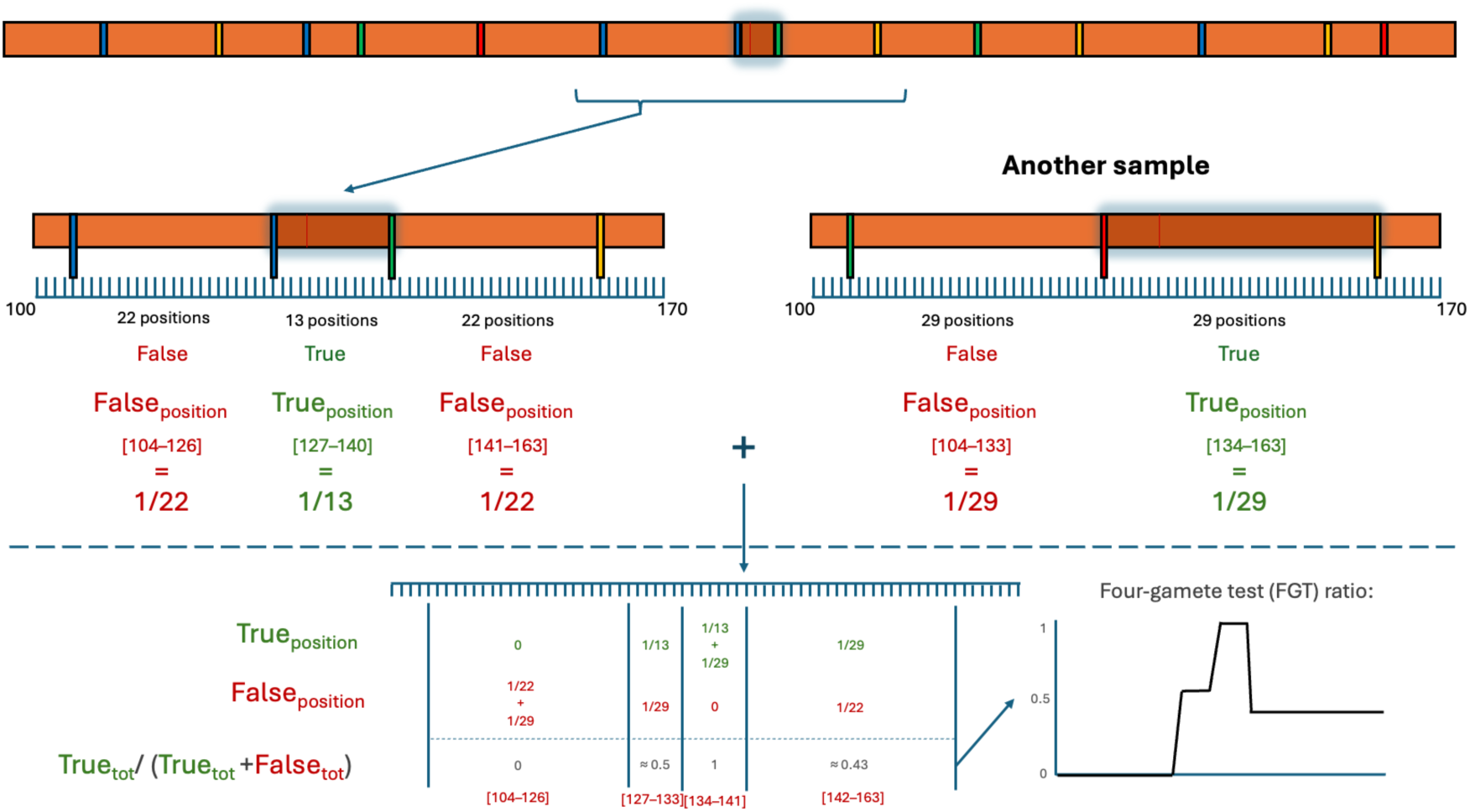
Converting discrete pair calls into a per-site FGT ratio profile. Each sample yields a set of position pairs classified as “True” (FGT satisfied) or “False” (FGT not satisfied). To obtain a genome-wide positional signal, each pair contributes uniformly across the genomic interval spanned by its two sites: a pair spanning (A-B) bases contributes a weight w=1/(B–A) to every covered position, adding to T(x) if “True” or to F(x) if “False”. Two per-position tracks, T(x) and F(x), which represent the total True and False support at nucleotide resolution, are produced through the summation of these contributions across all pairs. The per-site FGT ratio is then computed as R(x)=T(x)/(T(x)+F(x)), yielding a continuous positional profile that can be aggregated across multiple different SRA and visualized genome-wide.

To aid the interpretation of genome-wide profiles and to diagnose library and amplicon-driven artifacts, SRARec can additionally output two auxiliary read-distribution tracks. These include *(1)* the analysis coverage, defined as the per-nucleotide depth contributed only by reads that pass the analysis filters and that are, therefore, eligible to support variant linking and four-gamete comparisons, and *(2)* the read start/end density track computed by counting how many aligned reads start/end at each genomic coordinate. These counts are normalized to provide comparable positional profiles. While analysis coverage reflects how much usable evidence is available across the genome, read start/end density highlights clustering of alignment boundaries, as is often expected to occur under amplicon-based sequencing, and it helps flag regions where apparent signals may be influenced by non-uniform read boundaries and by position-specific error patterns that are common near the start/end positions of reads [82–84].

### The SRARec graphical user interface

Although large batches of SRA–NCBI IDs, FASTQ files, and BAM files can efficiently run on high-performance systems, we developed a user-friendly GUI optimized for macOS. The GUI allows users to select the data source (current version available only for BAM files) and adjust filtering and estimation parameters (i.e., minimum gamete frequency and support number of reads per gamete), and users can choose to ignore indels in the analysis. The results include an interactive visual summary consisting of the FGT ratio across the genome with optional annotation of the target viral genes drawn from the virus genome in its corresponding GenBank record. These outputs can be exported in different formats, including all raw information, a readable and organized description, or figures in interactive HTMLs or PNG. Download and more information about usage and capabilities is available at https://github.com/ldgonzalezvazquez/SRARec/.

### SRARec validation

We validated SRARec using simulated data based on three complementary evolutionary scenarios that include simulated sequences, real sequences upon which recombination events were simulated, and benchmark real datasets from previous studies.

We simulated mutations in the SARS-CoV-2 reference (NCBI NC_045512.2) to generate two sequences with different nucleotide diversity (π = 0.005, 0.01 and 0.05), insertion rates (0.0 and 0.0025) and deletion rates (0.0 and 0.0025). These sequences were recombined using random breakpoints to generate 2 homologous recombinant sequences. Next, we mutated the four sequences, further increasing nucleotide diversity (by π += 0.0 and 0.001), the insertion rate (0.0 and 0.00025) and the deletion rate (0.0 and 0.00025). Using the resulting sequences, we simulated FASTQ (pair end) files with the *NGSNGS* tool [85] at different coverages (200, 350, and 500) and read sizes (300, 500, and 1000), with a simulated sequencing quality of 100 (a parameter predefined in *NGSNGS*). Thus, we ran SRARec applying the implemented selection filters that include the minimum number of reads per polymorphism (5, 10, and 20), minimum number of reads per gamete (5, 10, and 20), and minimum minority gamete frequency (5%, 10%, and 20%).

After evaluating SRARec across this range of simulated conditions, we designed a second set of simulations to assess its performance under a more SARS-CoV-2-like evolutionary scenario, parameterized using a mutational profile reported for the SARS-CoV-2 Omicron variant of concern (VOC) [86]. Specifically, nucleotide diversity was estimated at 0.0022, whereas insertion and deletion rates were estimated at 0.00010033 and 0.00053507, respectively. Based on these estimates, we simulated combinations of nucleotide diversity values (π = 0.001, 0.0022, and 0.005), insertion rates (0 and 0.00010033), and deletion rates (0 and 0.00053507). Additional post-recombination variation was introduced by applying increments in nucleotide diversity (by π += 0 and 0.0003), and insertion and deletion rates (either 0 and 0.00005). Paired-end reads were subsequently generated using NGSNGS under the conditions described above. SRARec was then applied to the resulting simulated datasets using the same search parameters employed for the first set of simulations, except that the 20% minor-gamete-frequency threshold was excluded. This threshold was considered overly stringent because, under a four-gamete configuration, the maximum theoretical frequency of the least represented gamete is 25%, a value that would itself be unlikely under realistic conditions.

Next, to better approximate real SARS-CoV-2 evolutionary scenarios, we simulated recombination events with random breakpoint sites between real VOC sequences previously classified in [87] (mixture whole genome dataset composed of 50 sequences from each VOC). The procedure for categorizing detected recombination breakpoints was the same as previously described.

As an additional benchmark, we ran SRARec on the NCBI SRA entries that were used in two previously published analyses of intra-host SARS-CoV-2 recombination [47,51] carried out using different methodologies. To maximize detection power, for these simulations we relaxed the filters to five reads per polymorphism, and one read per gamete.

In [47], recombination was analysed using three FGT-based methods, analysing multiple sequence alignments of all reads flanking the S and N genes in a coinfected patient who was sequenced four times. Thus, we ran SRARec on all accessions deposited at SRA-NCBI for that study.

In [51], frequency imbalances and defining mutations of concordant variants with recombination were analysed. Of the 118 SRA-NCBI accessions extracted from the supplementary material of that article and claimed to be SARS-CoV-2, 23 (i.e., SRR12061247, SRR12859897 and SRR12825768) were not samples of that virus (verified in the NCBI) or were samples from private experiments and, therefore, had to be discarded. SRARec was run on the remaining entries.

### Applications of SRARec to the analysis of extensive SARS-CoV-2 and HIV-1 data

We retrieved accession identifiers for all NCBI SRA SARS-CoV-2 entries available as of September 4, 2025, and for HIV-1 SRA entries deposited by the PANGEA-HIV consortium under ENA accession PRJEB19239 [88]. Next, we ran SRARec using default parameters and the “slow” mode. For SARS-CoV-2, the analyses were performed by direct mapping to the reference (NC_045512.2). For HIV-1, we applied the high-divergence hybrid preprocessing strategy (*de novo* assembly of a unique reference for each analysed SRA accession followed by reference-guided refinement with *SPAdes/Shiver*, see above) and excluded indels from read-level haplotype comparisons. To report genome-wide profiles on a common coordinate system, we used the coordinate-mapping tables produced by *Shiver* to project per-sample results onto K03455.1 (HXB2) coordinates before aggregation and visualization. This produced results for 601,045 SARS-CoV-2 (Supplementary file 1, SC2 SRAs) and 4,999 HIV-1 SRA accessions.

To evaluate whether the genome-wide FGT ratio profiles were associated with biological features or sequencing-related biases, we compared the observed locations of peaks and valleys with a permutation-derived null distributions test (for additional information see Supplementary Material). The obtained FGT ratio was compared against three position technical tracks: sequencing coverage, reflecting the number of reads that entered in the analysis covering each genomic position; read start/end density, reflecting the local accumulation of read boundaries along the genome; and inter-polymorphisms interval coverage, reflecting the cumulative number of genomic positions lying between the two sites evaluated in all pairwise comparisons. For SARS-CoV-2, we additionally tested whether the FGT ratio profile overlapped with TRS-like sequence locations and with previously reported Sarbecovirus recombination profiles (recombination rate/rho, breakpoint distribution and breakpoint clustering significance) derived from analyses of the dataset constructed by De Klerk *et al.* [89] performed with RDP5 [12,13,90]. For HIV-1, we perform the same test using another dataset constructed by Simon-Loriere *et al.* [91].

For structural contextualization of selected genome regions, we generated RNA secondary-structure diagrams with VARNA (v3.93) [92] using SARS-CoV-2 genome-wide RNA structural information derived from [93]. Protein structure visualizations and residue-level coloring were produced with UCSF ChimeraX (v1.5), following standard ChimeraX workflows for structure rendering and annotation [94,95]. The used PDB accessions and structure sources were 7NNG for Helicase [96], 8GQC for the SUD region of nsp3 [97], 6YYT for RdRP [98] and 6XDC for ORF3a [99]. For Spike, we used 6VSB which represents the predominant prefusion conformation [100], with a receptor-binding domain in the up state. The regions of interest in protein M (amino acid sites 3-13) and the N5 loop within the Spike NTD (amino acid site 258) could not be physically represented because their coordinates were unresolved in the available structures.

## Results

The presented SRARec tool provides scalable read-level detection of intra-host recombination from large sequencing datasets. In particular, it accounts for minor variant information preserved in raw reads, rather than consensus sequences, to identify local haplotype incompatibilities consistent with FGT signals caused by recombination. By aggregating these read-level signals across samples and genomic positions, SRARec can be used to detect weak intra-patient recombination patterns at genome-wide scale.

### The validation tests indicated accurate recombination breakpoint detection in both simulated and real data

All possible filter combinations for each of the modes, “fast” and “slow”, were applied twice, resulting in 155,520 simulations. The analysis of the simulated data generated benchmarks with consistently high detection performance across coverage and read-length settings, with the expected trade-off that more stringent read-support and minor-gamete-frequency filters reduced false positives at the cost of some sensitivity. When we simulated the diversity and insertion and deletion rate based on those reported for Omicron [86], insertions had little effect on performance, whereas deletions moderately reduced detection, likely because they interfered with read mapping.

Overall, the detection accuracy improved with higher diversity and longer reads, whereas indels increased false positives by degrading mapping accuracy (Supplementary file 1, Slow and Fast modes).

Regarding the 9,000 simulations based on real SARS-CoV-2 variants, they yielded a more conservative detection rate but maintained a low false-positive count (n=2) (Supplementary file 1) (González-Vázquez 2025).

Finally, when validating the software on empirical datasets studied in previous works, breakpoint detections obtained with SRARec clustered near the recombination sites identified in those studies, although they were not completely identical. In addition, SRARec identified complementary recombination signals beyond those captured in the original analyses, likely because both published studies encountered regional limitations by exploring a relatively small subset of genome regions with enough genetic diversity to detect recombination, or only the polymorphisms that defined the two variant lineages involved in the mixed infections (Supplementary file 1) (Wertheim 2022 and Pipek 2024).

### Genome-wide intra-host recombination signatures differ between HIV-1 and SARS-CoV-2

We applied SRARec to study genome-wide recombination profiles from large publicly available short-read datasets, specifically 601,045 SARS-CoV-2 SRAs and 4,999 HIV-1 SRAs. We found markedly different genome-wide distributions of detectable intra-host recombination signals in the two viruses (Figures 7 and 8; see Supplementary files 2 and 3 for interactive visualization). Specifically, HIV-1 showed FGT ratios of between 0.45 and 0.72 across 95% of sites in the genome, with sustained high FGT ratios over long genomic stretches rather than being confined to a small number of narrow “hotspot” sites or regions (Figure 7). In SARS-CoV-2, the genome-wide profile showed a predominantly low background FGT ratio, punctuated by comparatively infrequent, localized elevations (Figure 8). This difference is also reflected in the dynamic range of the FGT ratios, with a higher and more pervasive read-level signal detected in HIV-1. However, this should not be interpreted as evidence of a directly comparable higher recombination rate, because polymorphisms were substantially more prevalent across the HIV-1 SRA accessions, with 95.62% of genomic positions containing polymorphisms detected in at least 1% of HIV-1 SRAs, compared with only 0.22% of genomic positions in the SARS-CoV-2 SRAs. This provided substantially more opportunities for recombination detection in HIV-1. Note that these proportions include only polymorphisms that entered the SRARec analysis, namely those covered by reads spanning at least two polymorphic sites.

**Figure 7.**
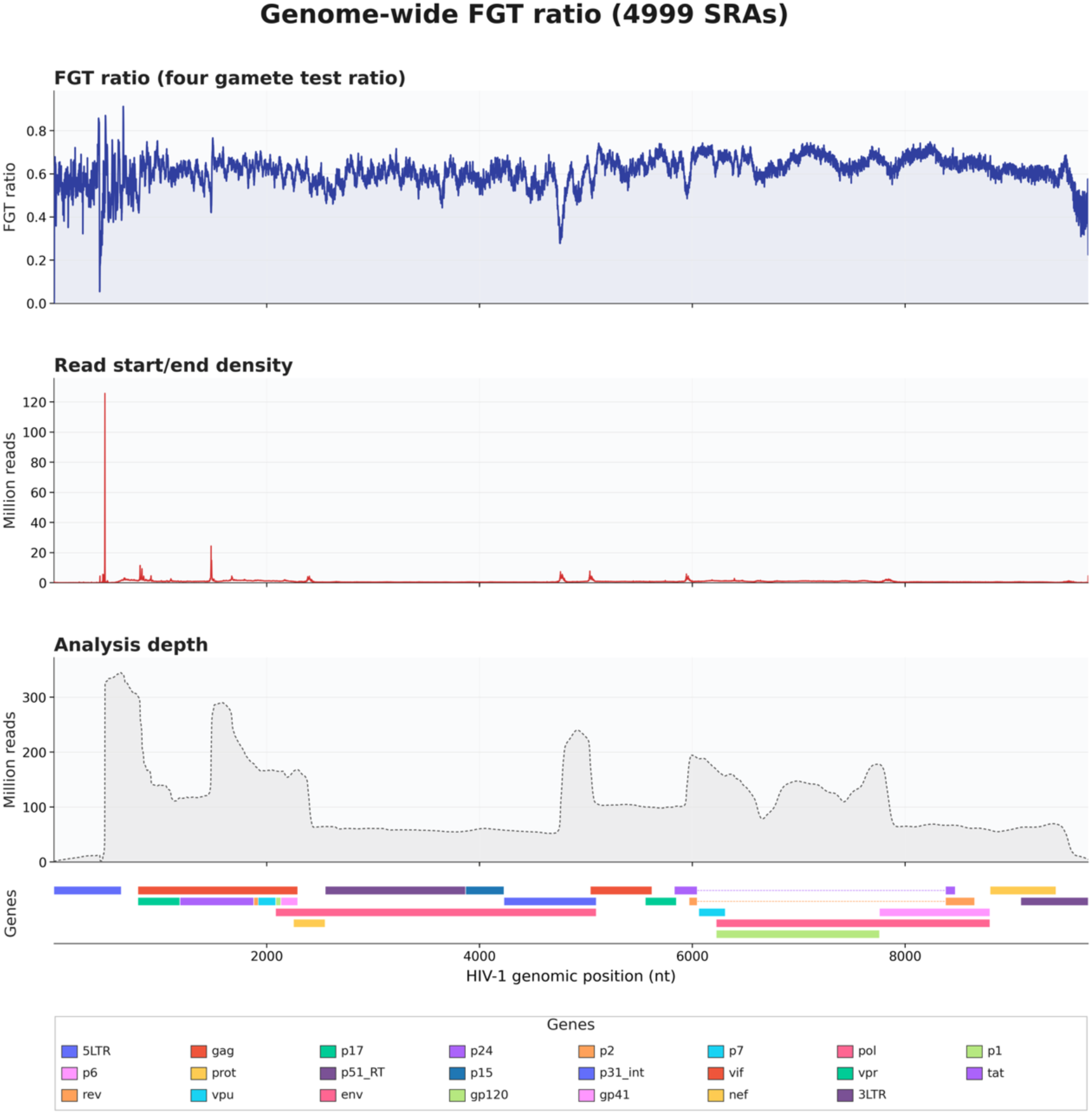
Aggregated intra-host genomic recombination profile across 4,999 Human Immunodeficiency Virus type 1 (HIV-1) Sequence Read Archive (SRA) accessions. The plot displays the recombination landscape across the HIV-1 genome, based on the analysis of 4,999 SRA datasets. As indicated in the figure, the top panel (blue) represents the four gamete test (FGT) ratio, the middle panel (red) shows the density of read starts and ends (in millions of reads), and the bottom panel (grey) indicates the analysis depth (in millions of reads) (see Methods for more detail). The FGT ratio is calculated across all SRAs as the ratio of positive tests to total tests, where each site per sample is evaluated as a binary (positive/negative) value, as detailed in the Methods section. A legend of coding regions is shown at the bottom, drawn according to the National Center for Biotechnology Information (NCBI) K03455.1 reference. See Supplementary file 2 for interactive visualization.

**Figure 8.**
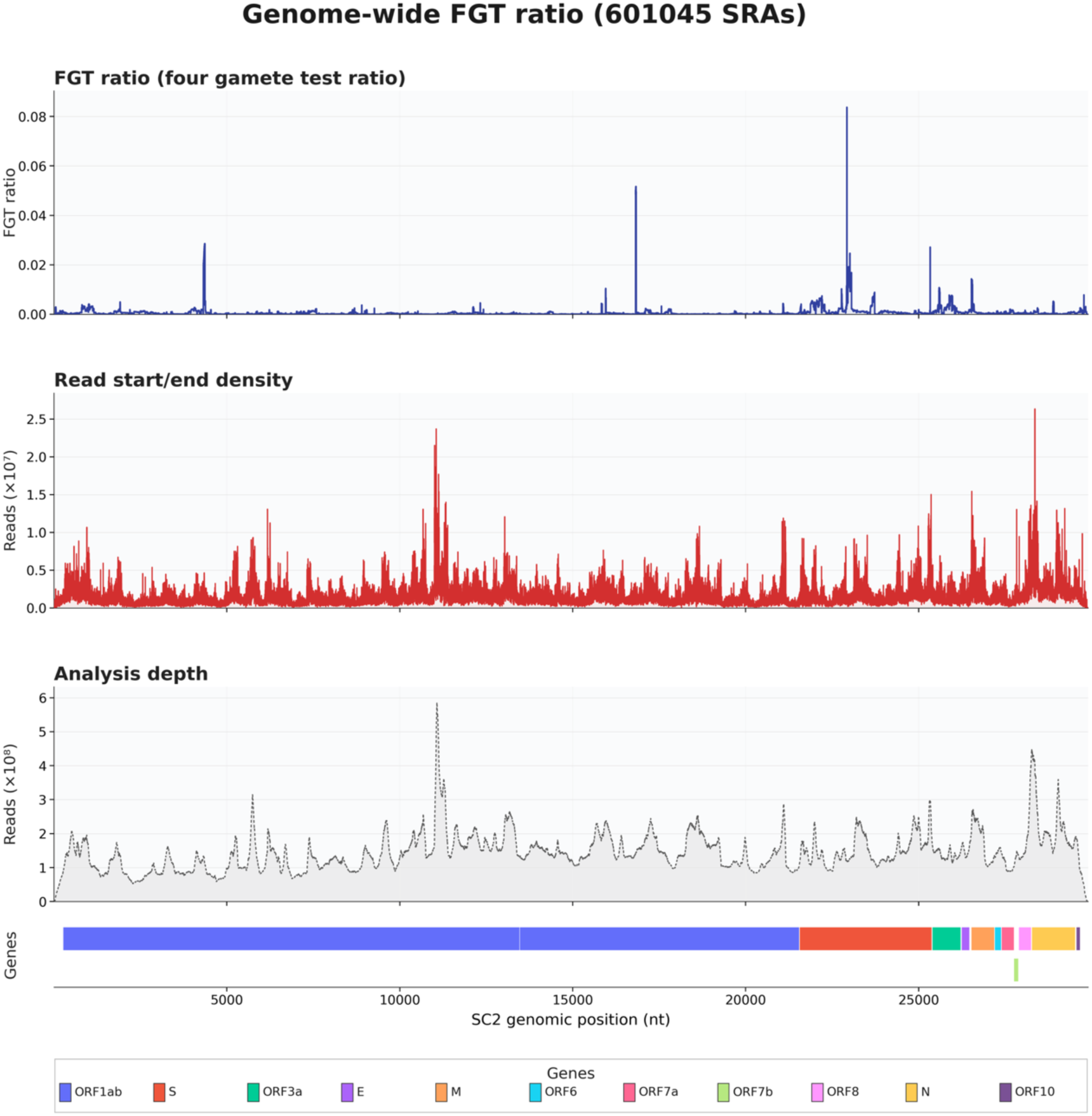
Aggregated intra-host recombination profile across 601,045 Severe Acute Respiratory Syndrome Coronavirus 2 (SARS-CoV-2) Sequence Read Archive (SRA) accessions. The plot illustrates the recombination landscape across the SARS-CoV-2 genome, generated from the screening of 601,045 SRA datasets. As shown in the figure, the top panel (blue) represents the four gamete test (FGT) ratio, the middle panel (red) denotes the density of read starts and ends (×10⁷ reads), and the bottom panel (grey) displays the analysis depth (×10⁸ reads) (see Methods for more detail). The FGT ratio is calculated across all SRAs as the number of positive tests divided by the total tests, where each site per sample is evaluated as a binary (positive/negative) value, as explained in the Methods section. A legend of coding regions is shown at the bottom, drawn according to the National Center for Biotechnology Information (NCBI) NC_045512.2 reference. See Supplementary file 3 for interactive visualization.

### Pervasive genome-wide intra-host recombination profile in HIV-1

The screening of HIV-1 SRAs revealed a pervasive, elevated, and sustained recombination background across the genome (Figure 7). The baseline FGT ratio consistently fluctuated within a high dynamic range between 0.00 and 0.91. The recombination profile exhibited distinct localized deviations from this high baseline. For instance, at the 5’ terminus of the genome, the landscape displayed prominent fluctuations, showing sharp rises and dips in the FGT ratio throughout the 5’ LTR region. Moving downstream through the structural and enzymatic coding regions, the ratio remains relatively stable until reaching the transition at the end of the polymerase gene. Specifically, within the region encoding the integrase p31, the profile displays a sudden, highly pronounced, and localized dip where the FGT ratio drops sharply to approximately 0.3. After returning to its elevated baseline throughout the accessory and envelope genes, a final deviation occurs at the 3’ end of the genome, where we found a decline in the recombination signal observed across the 3’ LTR region.

To test whether the HIV-1 FGT ratio profile could be explained by methodological related heterogeneity, we compared it against inter-polymorphisms interval coverage, sequencing coverage and read start/end density (Figures S1-S3; Supplementary Material). A permutation analysis explained in Supplementary Material showed that none of the three signals were statistically related to our recombination profile (FGT ratio; p-values of 0.404, 0.788 and 0.595, respectively). To test whether our signal mimicked known HIV-1 recombination patterns, we applied the same test against three recombination profiles calculated with RDP5 on a dataset published in [91] (see methods). We found associations with the population recombination rate (ρ) and the distribution of breakpoints (p-values 0.046 and 0.005, respectively), but not with clustering of breakpoints per genomic region (p-value 0.469). These results indicated that our signal was not the result of a technical imbalance between positions and recovered signals also recoverable from population consensus studies (Figures S4-S6; Supplementary Material).

### SARS-CoV-2 has a low genome-wide recombination background with some local hotspot regions

In contrast to the pervasive recombination signal detected in HIV-1, the SARS-CoV-2 FGT ratio showed discrete recombination hotspots (FGT ratio > 0.01) in a limited number of genomic regions (Figure 8). The strongest recombination hotspot signal lies within the Spike coding region, spanning a short interval in the receptor-binding region (nt 22927-23062, which corresponds to spike codons 455 to 500, Figure 8). A second single-position recombination Spike-associated hotspot was observed at a different site within the Spike coding region (nt 22336, which corresponds to spike codon 258, Figure 8) accompanied by a narrower focal signal in the core of the Receptor-Binding Domain (RBD) encoding region of the spike gene (nt 22770-22774, which corresponds to spike codons 403-404, Figures 8) and a broader, sub-threshold increase extending across the region of the spike gene encoding the N-terminal domain (nt 21875-22286, which corresponds to spike codons 105-242, Figure 8). Outside the Spike region, additional recombination hotspots were detected within ORF1ab, including an interval in nsp3 (nt 4323-4366, which corresponds to ORF1ab codons 1353-1367, Figure 8), a narrow segment in nsp12/RdRp (nt 15951-15964, which corresponds to ORF1ab codons 5229-5234, Figure 8), and a short region in nsp13/helicase (nt 16821-16835, which corresponds to ORF1ab codons 5519-5524, Figure 8). Finally, in the 3’ structural and accessory region, localized increases are detected within the ORF3a transmembrane encoding region (nt 25592-25607, which corresponds to ORF3a codons 67-72, Figure 8) and at the beginning of the M coding region (nt 26529-26560, which corresponds to M codons 3-13, Figure 8). Together, these results indicate that the aggregated detected recombination signal across 601,045 SARS-CoV-2 intra-host datasets is not uniformly distributed throughout the genome but is largely concentrated to certain narrow genomic intervals.

To provide structural context to the detected hotspot intervals, we mapped each region onto available protein structures (Figures 9, 10 and 11). In the protein renderings, those segments correspond to localized patches within the three-dimensional folds. For Spike, the RBM-associated interval (codons 455-500, Figure 9B) and the secondary focal signal in the RBD (codons 403-404, Figure 9A) form surface patches in the region of the spike protein that interacts with the ACE2 receptor of target cells. Similarly, localized patches were visible for the ORF1ab hotspots mapped to nsp3 (codons 1353-1367; Figure 10A) and nsp13/helicase (codons 5519-5524; Figure 10C), as well as for the focal signal within the ORF3a transmembrane region (codons 67-72; Figure 9C). In contrast, the RdRp-associated hotspot (codons 5229-5234; Figure 10B) was less apparent as a surface patch when displayed in the context of the full polymerase assembly, where the detected region is positioned more internally.

**Figure 9.**
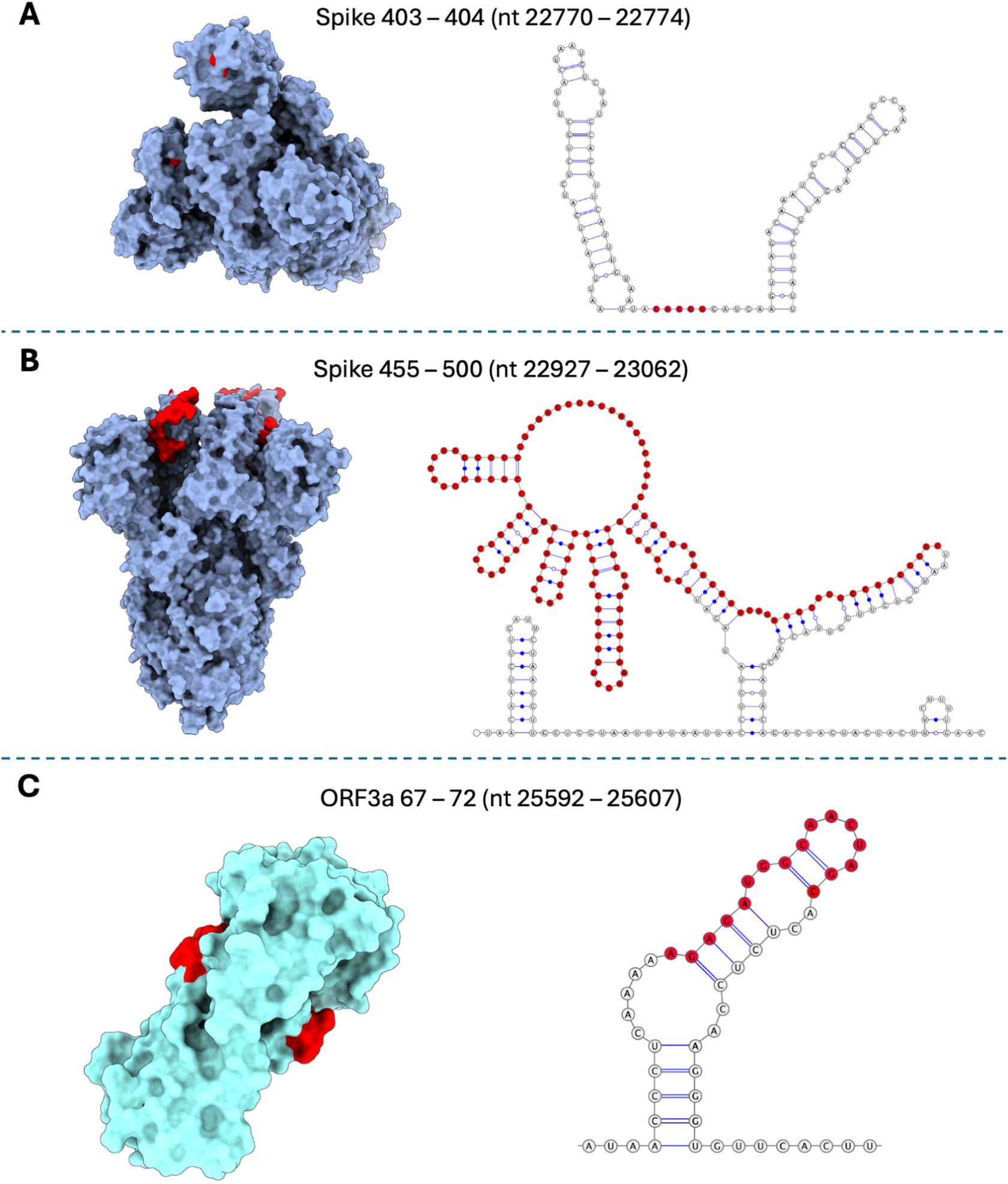
Protein and RNA structural mapping of SARS-CoV-2 Spike glycoprotein and ORF3a recombination hotspots. Three-dimensional protein representations and their corresponding RNA secondary structure models, with primary recombination signals highlighted in red. Spike glycoprotein homotrimer illustrating: (A) the focal recombination signal (nt 22770-22774 corresponding to codons 403-404) located in the Receptor-Binding Domain (RBD), paired with its corresponding RNA sequence folding; and (B) the major recombination hotspot (nt 22927-23062 corresponding to codons 455-500) mapped onto the exposed Receptor Binding Motif (RBM), shown in the Spike conformation with one RBD up and two RBDs down [100], alongside the local RNA secondary structure of this interval. (C) ORF3a accessory protein that illustrates the recombination signal (nt 25592-25607 corresponding to codons 67-72) located within the transmembrane region of the ORF3a accessory protein, shown together with its respective RNA sequence folding.

**Figure 10.**
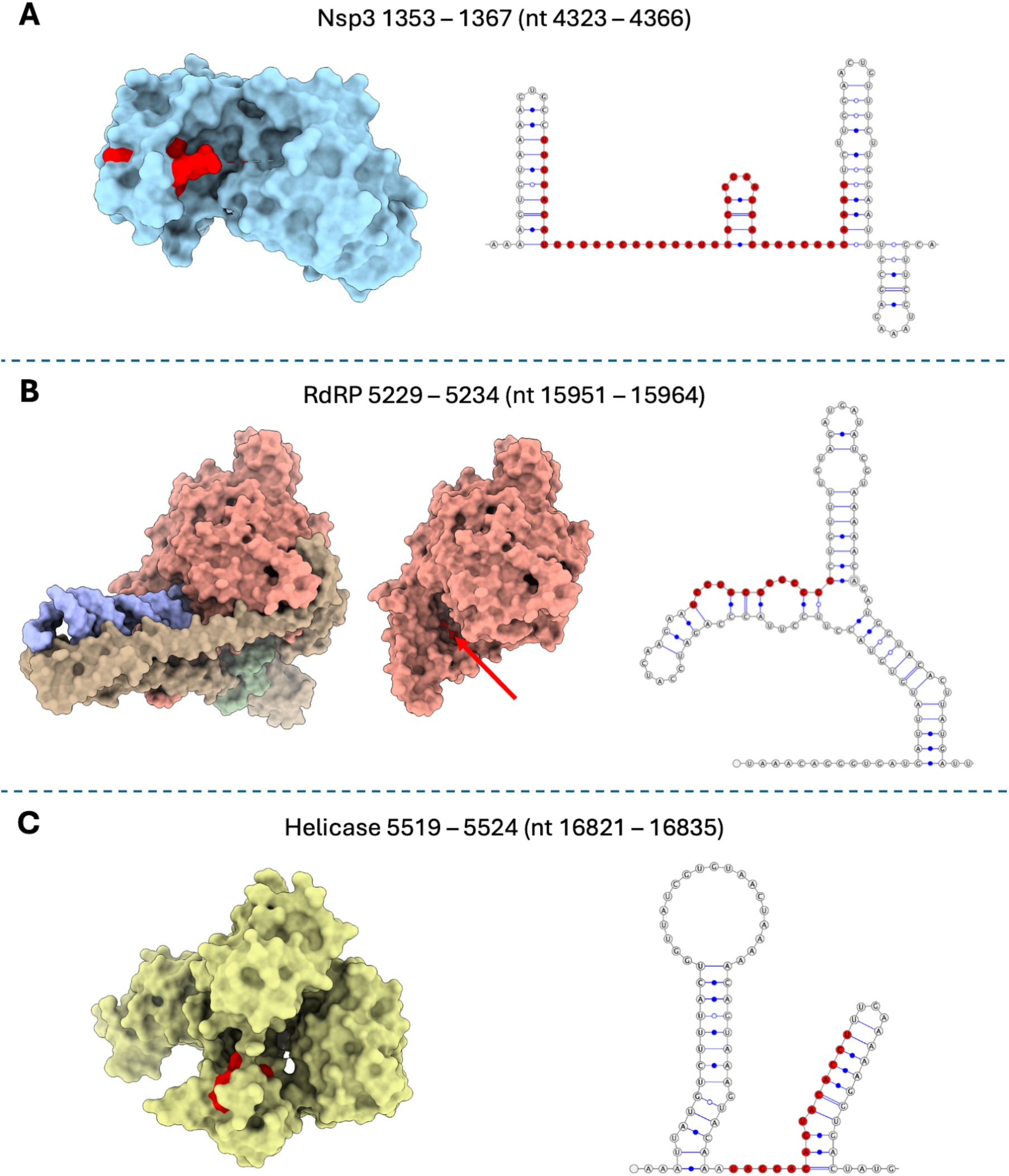
Structural and RNA mapping of SARS-CoV-2 ORF1ab recombination hotspots. Three-dimensional protein representations and their corresponding RNA secondary structure models, with primary recombination signals highlighted in red. (A) SARS-Unique Domain (SUD) of nsp3, highlighting the surface-exposed nature of the signal (nt 4323-4366 corresponding to codons 1353-1367), shown together with its respective RNA secondary structure. (B) RNA-dependent RNA polymerase (RdRp, nsp12) complex showing the deep internal localization of the recombination signal (nt 15951-15964 corresponding to codons 5229-5234), paired with its corresponding RNA sequence folding into a stable stem-loop. (C) Helicase (nsp13) structure, with the recombination signal (nt 16821-16835 corresponding to codons 5519-5524) falling within the flexible 1B-domain loops, shown together with its respective RNA secondary structure.

**Figure 11.**
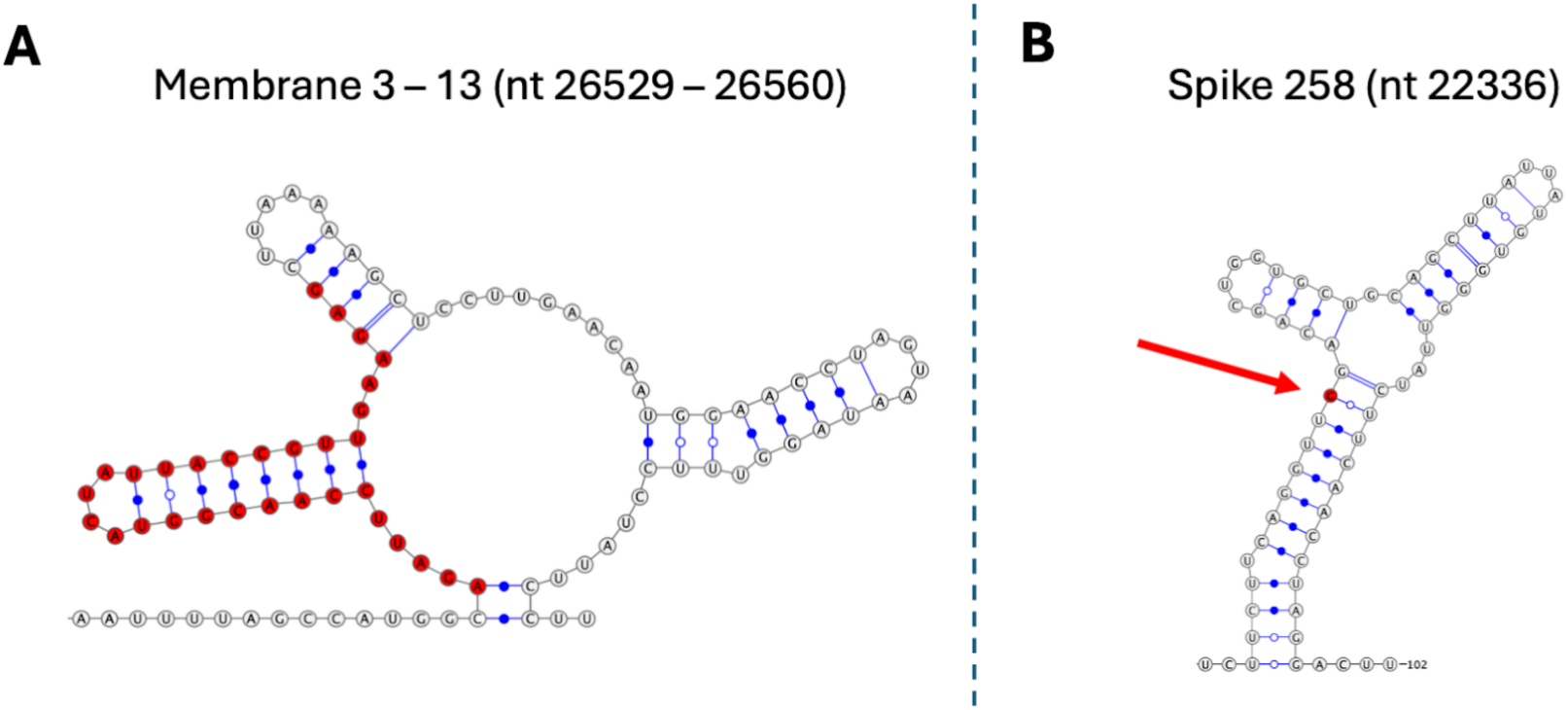
RNA structural mapping of recombination hotspots in SARS-CoV-2 Membrane and Spike N-terminal domains. RNA secondary structure models corresponding to genomic intervals with unresolved three-dimensional protein coordinates, with primary recombination signals highlighted in red. (A) Local RNA secondary structure of the localized recombination signal detected at the beginning of the Membrane protein coding region (nt 26529-26560 corresponding to codons 3-13). (B) Local RNA sequence folding associated with the focal hotspot (nt 22336 corresponding to codon 528) located within the N-terminal domain of the Spike protein.

At the RNA level, we mapped the most intense recombination signals to corresponding segments of the SARS-CoV-2 RNA secondary-structure model (Figures 9, 10 and 11). Across loci, including the Spike-associated signals (nt 22927–23062 and nt 22336 corresponding to codons 455-500 and 258 respectively, Figures 9B and 11B), the ORF1ab hotspots (nsp3, RdRp and helicase, Figure 10), the signal at the beginning of M (nt 26529– 26560 corresponding to codons 3-13, Figure 11A) and in transmembrane encoding region of ORF3a (25592–25607 corresponding to codons 67-72, Figure 9C), the recombination hotspot nucleotide region fell within highly branching local secondary structural elements within the RNA structural model.

To test whether the SARS-CoV-2 FGT ratio profile could be explained by methodological related heterogeneity, we compared it against inter-polymorphisms interval coverage, sequencing coverage and read start/end density applying the same permutation test that we performed in the case of HIV-1 (Figures S7-S9; Supplementary Material). We found that none of the three signals were statistically related to our recombination profile (p-values of 0.247, 0.629 and 0.488, respectively). To test whether our signal mimicked known biological or evolutionary patterns, we also applied the same test against TRS-like similarity regions and two Sarbecovirus recombination profiles (breakpoint distribution and recombination rate) provided by De Klerk *et al.* [89] (Figures S10-S12; Supplementary Material). We found no significant associations with TRS-like regions (p-value 0.597), the breakpoint distribution profile (p-value 0.548) or the recombination rate (p-value 0.405).

These results indicated that our signal was not the result of a technical imbalance between positions and that the detected recombination hotspots did not simply recapitulate population-level Sarbecovirus or TRS-driven patterns.

## Discussion

By evaluating minor variant haplotypes directly from raw sequencing reads, SRARec captures intra-host recombination dynamics that are otherwise masked by consensus-sequence based recombination analyses. Our large-scale screening revealed distinct genome-wide distributions of detectable read-level recombination signals in HIV-1 and SARS-CoV-2. HIV-1 showed a broadly distributed and relatively uniform signal across 4,999 accessions, whereas SARS-CoV-2 showed a low background punctuated by discrete localized peaks across 601,045 infections. However, a substantial part of this contrast is likely driven by differences in the opportunities for recombination detection rather than by directly comparable differences in the absolute recombination rates of the two viruses. The greater within-host diversity of HIV-1 provides many more informative polymorphic site pairs per SRA accession, increasing the probability that recombinant haplotypes will be detectable by the FGT. By contrast, the lower within-host diversity typically observed in SARS-CoV-2, together with shorter infection durations and narrow transmission bottlenecks, greatly limits detection opportunities. These profiles therefore demonstrate the capacity of read-level analysis to characterize the spatial distribution of detectable recombination signals within divergent RNA viruses, but they should not be interpreted as a direct quantitative comparison of their underlying recombination rates.

### Methodological resolution and technical robustness of read-level FGT screening

The application of the four-gamete test (FGT) directly to raw sequencing reads provides a lower bound of the true intra-host recombination frequency. Because recombination detection requires physical linkage within individual reads, this local resolution cannot capture recombination events between polymorphisms that exceed the sequencing read length, and sensitivity necessarily drops if parental genomes are nearly identical or lack informative read co-coverage. However, by completely bypassing consensus sequence assemblies, this approach successfully uncovers genuine, low-frequency recombinant haplotypes closer to their biological origin before they are masked or artificially generated by assembly algorithms. Furthermore, our analysis demonstrates that this read-level signal is robust against protocol-driven biases. Although amplicon-based sequencing workflows (including widely used schemes such as ARTIC [101,102]) introduce severe positional non-uniformity in sequencing coverage and read boundary density [83,84], permutation-based analyses showed no significant association between these technical profiles and the FGT ratio landscapes in either virus. The lack of statistical enrichment for different analytical measures in HIV-1 and SARS-CoV-2 analyses showed genuine biological footprints rather than artifacts of library preparation or sequencing topography.

### Interpretation of the diffused and pervasive HIV-1 recombination landscape

The analyses of HIV-1 reveal a broadly elevated and sustained recombination signal across most of the genome (Figure 7), consistent with recombination acting as a pervasive evolutionary force within individual infections. This pattern aligns with known HIV-1 recombination mechanisms, where the encapsidation of two RNA genomes per virion facilitates repeated template switching during reverse transcription, with recombination occurring at a frequency comparable to point mutation [103]. Together with high within-host diversity and the frequent coexistence of multiple genetically distinct lineages over prolonged infections [67,104], these properties enrich the pool of informative polymorphisms and linked haplotypes, rendering read-level recombination signals significantly more detectable and broadly distributed than in SARS-CoV-2.

Despite this high baseline, the HIV-1 FGT ratio profile exhibits localized fluctuations that likely reflect a heterogeneous landscape of hotspots and coldspots. In the 5′ leader/gag region, these variations occur where complex RNA secondary structures guide dimerization, genome packaging, and reverse-transcription initiation [105,106]. Outside the 5′ region, the signal remains broadly elevated throughout the accessory and envelope gene regions. In the envelope gene, intense immune selection maintains high local diversity [107,108], providing abundant informative variation for detection. However, this diversity interacts with strict selective constraints. In particular, RNA secondary structure could influence hotspots in gp120, while purifying selection against dysfunctional recombinants could shape observable breakpoint distributions even within individual infections [91,109,110].

Functionally, this pervasive recombination provides a powerful mechanism for exploring sequence space by reshuffling standing variation [111]. It can also accelerate adaptation by combining beneficial alleles from distinct genetic backgrounds into a single genome while simultaneously uncoupling them from linked deleterious mutations. Clinically, such dynamics facilitate the rapid assembly of multi-mutation genotypes including linked drug-resistance mutations and immune-escape substitutions [111], thereby driving therapeutic escape and contributing to the extensive mosaicism characteristic of circulating HIV-1 lineages [103].

### Interpretation of the sparse and focal SARS-CoV-2 recombination landscape

The sparse distribution of the SARS-CoV-2 FGT ratio spikes suggests that while observable intra-host recombination is likely a rare event under typical infection dynamics, these signals tend to recur at non-random genomic regions (Figure 8). When contextualized with protein structures and RNA models (Figures 9, 10, and 11), these hotspots can be interpreted through three main hypothetical mechanisms:

*(i)* Potential adaptive shuffling at the host-immune interface. In the Spike gene, the primary signals cluster at sites encoding flexible, solvent-exposed loops of the RBM (codons 455-500) and NTD (codon 258) [100,112], which constitute the main interfaces for immune pressure and cell entry [113,114]. Recombination within these domains (particularly around residues 455–500 [115,116] and the RBD core [117]) could allow the virus to rapidly assemble multi-mutation genotypes that might optimize receptor affinity or facilitate antibody escape [46]. The documented tolerance of NTD loops to structural changes [114,118] and the emergence of “FLip” mutations [119,120] support the hypothesis that these domains could act as relatively exchangeable modules capable of generating advantageous recombinant variants.
*(ii)* Locus-specific transcription and domain modularity. Although TRS elements do not show a genome-wide correlation with the locations of detectable recombination breakpoint peaks, the signals detected near the beginning of the M (codons 3-13) and ORF3a (codons 67-72) genes remain compatible with localized, transcript-guided template switching [37,40]. This pattern suggests the potential existence of post-TRS “landing zones” where the polymerase might stabilize its hybridization [37,121]. Such chimeric events might only be viable in specific regions. For instance, the M gene signal maps to the exposed N-terminal ectodomain [122], whereas events further downstream in the rigid transmembrane domains could result in unviable virions [123]. This structural tolerance might also explain the signals in the surface-exposed nsp3 [97] and the transmembrane region of ORF3a [124], where domain swapping could occur without fully disrupting the polyprotein core. However, this modular tolerance remains highly hypothetical, as the accumulation of sufficient genetic diversity within a single individual infection to cause severe recombination-induced protein misfolding problems is generally unlikely, though such structural conflicts could become more probable during coinfections involving highly divergent variants.
*(iii)* Mechanical pausing in the replication core. In contrast to surface-exposed regions, the internal recombination breakpoint clustering within the RdRp (codons 5229-5234, Figure 10B) maps to stable RNA stem-loops that are thought to induce polymerase pausing, which could mechanically trigger template switching [125]. Similarly, the helicase signal (codons 5519-5524) falls within codons encoding flexible 1B-domain loops [96] targeted by CD8+ T-cells [126,127], which could tentatively offer a mechanical route to evade cellular immunity while preserving essential replication mechanics. Importantly, in regions where genetic diversity is minimal, it is theoretically easier for the transferred strand to hybridize properly during template switching [2,37]. Therefore, the high sequence conservation of the RdRp core [128] could paradoxically act as a major structural facilitator of recombination at this specific locus.

Overall, the distribution of these SARS-CoV-2 FGT ratio increases suggests that detectable intra-host recombination is likely gated by both structural viability and adaptive utility. Rather than random accumulation, these localized signals seem to cluster where the viral protein architecture appears to tolerate structural changes, in exposed loops under selective pressure (Spike), stabilizing transcription zones (M, ORF3a), or mechanical pause sites within the replication machinery (RdRp, Helicase).

### Surveillance implications, limitations, and diagnostic scope

By shifting the analytical focus from consensus sequences to raw read-level sequences, SRARec operates upstream of traditional recombination analysis methods, acting as an extra layer that uncovers cryptic intra-host diversity before it is altered by selective filtering or masked by consensus assembly artifacts. However, the analytical resolution of this screening is inherently gated by read linkage and sequence diversity. It requires that informative polymorphic positions are simultaneously co-covered by individual reads within sufficiently divergent parental backgrounds. Consequently, sensitivity varies with read length and library architecture and estimates at highly selected loci can tentatively be influenced by alignment ambiguity or homoplasy.

Despite these intrinsic constraints, bypassing consensus sequence generation remains essential for capturing the raw material of viral evolution and establishing a foundation for proactive genomic surveillance. For instance, the reproducible SARS-CoV-2 hotspot intervals that we detected define a highly constrained set of targets ideal for focused verification (Figure 8). This is particularly critical in the Spike gene, where intense antibody- and vaccine-driven selection concentrates [113], meaning that recombination could provide a rapid evolutionary shortcut to shuffle multi-mutation genotypes across different segregating backgrounds (as is observed in the circulating recombinant SARS-CoV-2 lineages like XBB [129,130]) rather than relying on slow, sequential mutation accumulation. The detection of these localized recombination hotspots for functional and epidemiological follow-up could significantly accelerate the early identification of emerging viral variants of potential public health relevance.

## Code availability

SRARec, including source code, GUI, and documentation, is publicly available from GitHub at https://github.com/ldgonzalezvazquez/SRARec and has been permanently archived in Zenodo at doi: 10.5281/zenodo.21699679. SRARec is distributed under the MIT license, which allows its free use, modification, and adaptation to study any data. Scripts used to run the simulations are also publicly available from GitHub at https://github.com/ldgonzalezvazquez/SRARec/tree/SRARec_simulations.

## Data availability

The raw sequence datasets analysed (SARS-CoV-2 and HIV-1 SRAs) are publicly available from the NCBI SRA database. The individual identifiers are provided in Supplementary file 1. PANGEA-HIV consortium datasets are deposited in ENA accession PRJEB19239 [88]. Simulation results are available in Supplementary file 1.

## Supplementary material

A main Supplementary Material together with Supplementary Files 1, 2, and 3 are available at the journal online.

## Funding

L.D.G.-V. was funded by a fellowship from Xunta de Galicia ED481A-2023/089, programa de axudas á etapa predoutoral da Xunta de Galicia (Consellería de Cultura, Educación, Formación Profesional e Universidades) cofinanciado pola Unión Europea no marco do Programa FSE+ Galicia 2021-2027. M.A. was initially supported by the Grant CNS2023-144363 funded by MICIU/AEI/10.13039/501100011033 and by European Union NextGenerationEU/PRTR, and finally supported by the Grant PID2023-151032NB-C22 funded by MICIU/AEI/10.13039/501100011033 and by FEDER, UE. Funding for open access charge: Universidade de Vigo/CISUG.

## Acknowledgements

We thank Centro de Supercomputación de Galicia (CESGA) for the computer resources.

## Conflicts of Interest

The authors declare no conflict of interest.

## Contributions

L.D.G.-V. (Conceptualization [equal], Formal analysis [lead], Funding acquisition [equal], Investigation [lead], Resources [lead], Validation [equal], Visualization [lead], Writing – original draft [lead], Writing – review & editing [lead]); P.I.-R. (Resources [supporting], Validation [equal], Visualization [supporting], Writing – review & editing [supporting]); M.A. (Funding acquisition [equal], Visualization [supporting], Writing – review & editing [supporting]); D.P.M. (Funding acquisition [supporting], Conceptualization [equal], Formal analysis [supporting], Investigation [supporting], Resources [supporting], Validation [supporting], Visualization [supporting], Writing – review & editing [supporting]).

## Notes

### Competing Interest Statement

The authors have declared no competing interest.

