## Supplementary Material for "SRARec: A program for detecting recombination in sequencing reads and its application to uncover recombination patterns in SARS-CoV-2 and HIV-1"

To test whether FGT ratio profiles were associated with genomic and biological features, or potentially affected by technical and methodological biases, we applied a permutation-based peak-enrichment analysis to our SARS-CoV-2 and HIV-1 results.

In this context, we mapped genome-wide comparison tracks corresponding to positional signals or annotations onto viral genome coordinates and analysed them against the FGT ratio profile. We initially focused on three technical tracks: sequencing coverage, read begin/end density and inter-polymorphisms interval coverage (Pair span). These tracks captured different aspects of the sequencing and comparison structure: local read depth, clustering of read starts and ends, and the cumulative number of times each genomic position was located between the two polymorphic sites of the pairs evaluated by SRARec.

For each comparison track, we classified genomic positions as peaks or valleys according to the values of that track. For SARS-CoV-2, we first used a fixed FGT ratio threshold of 0.01 to calculate the proportion of genomic positions exceeding this value. We then classified the same proportion of the highest-valued positions as peaks in the corresponding comparison track. For HIV-1, we defined peak positions using a fixed upper 5% cutoff.

After the initial peak/valley classification, we grouped consecutive peak positions into continuous blocks. We merged peak blocks separated by valleys shorter than 100 nucleotides into single continuous regions by filling the intervening valley. For each virus, we then derived a symmetric window size from the FGT ratio profile by applying the same procedure and calculating the median length across merged blocks, yielding 17 nt for SARS-CoV-2 and 119 nt for HIV-1. We applied this window to all FGT-anchored peak/valley masks we used in permutations and figures and only after raw thresholding and 100-nt peak merging. At each genomic position, we evaluated a centered window of the specified length and assigned peak status if we had already classified any position within that window as a peak (rolling maximum over the binary mask). In practice, this operation expands peak blocks outward in both directions, so narrow threshold-defined peaks become wider contiguous peak regions while valleys outside this expansion remain valleys. We always computed thresholds on raw signals; the window did not recalculate them but only modified the final binary P/V mask we used for enrichment testing and figure shading (see below).

For SARS-CoV-2, we also performed two additional exploratory comparisons. First, we tested whether the FGT ratio signal was enriched within TRS-like nucleotide regions. We defined TRS-like positions using a motif-similarity threshold allowing up to one mismatch. Second, we compared the SARS-CoV-2 FGT ratio profile with external Sarbecovirus recombination-breakpoint information obtained with RDP5 [1-3] from a dataset constructed by De Klerk et al. [4]. We converted these data into genome-wide comparison tracks (see methods) and analysed them using the same permutation test.

For HIV-1, we performed an additional comparison with recombination information obtained with RDP5 from the dataset constructed by Simon-Loriere *et al.* [5]. We included these analyses to evaluate whether the HIV-1 FGT ratio profile recapitulated independent recombination patterns inferred from previously analysed HIV-1 recombinant consensus sequence datasets. Specifically, we compared the FGT ratio profile with recombination-rate estimates, breakpoint-distribution signal and breakpoint-clustering significance (see methods).

For comparisons with recombination-related profiles exported from RDP5, we mapped external tracks onto the reference coordinate system we used in our analyses (SARS-CoV-2 NC_045512.2 for Sarbecovirus and HIV-1 HXB2-like coordinates for HIV-1). Because corresponding ORFs often differed in length between the external alignment and our reference genome, we adjusted coordinates ORF by ORF rather than by whole-genome linear scaling. For each shared ORF, we first expressed an external position relative to that ORF and then mapped it linearly onto the matching reference ORF interval. We transferred the associated signal value accordingly, using fractional weights when the mapped position fell between integer nucleotide sites. We did not infer or pad ORFs present in only one dataset. This procedure compresses or expands each external ORF onto its reference interval while preserving the relative signal distribution within that ORF. In figures, the upper panel shows the external track in native RDP5 alignment coordinates alongside the FGT ratio profile in reference coordinates, whereas the lower panel shows both signals after ORF-wise adjustment on the reference genome; we performed permutation tests (see below) for external RDP tracks on these adjusted, mapped profiles.

For each comparison, we first converted the tested genome-wide track into a binary peak/valley mask. We then calculated the observed enrichment statistic by multiplying the FGT ratio value at each genomic position by the binary peak/valley state of the tested track at that same position, and summing this product across all positions:

$$S_{obs} = \sum_{i} (R_i M_i)$$

where $R_i$ is the FGT ratio at genomic position $i$, and $M_i$ is the binary peak/valley state of the tested genome-wide track at that position:

$$\mathrm{M} = \begin{cases} 1, if\ position\ i\ belongs\ to\ a\ peak\ region\ of\ the\ tested\ track \\ 0, if\ position\ i\ belongs\ to\ a\ valley\ region\ of\ the\ tested\ track \end{cases}$$

Depending on the comparison, the tested track corresponded to sequencing coverage, read start/end density, inter-polymorphisms comparison coverage, TRS-like motif similarity, or external recombination-related features derived from Sarbecovirus and HIV-1 RDP5 analyses. Thus, $S_{obs}$ represents the total FGT ratio signal accumulated within the peak regions of the tested track.

We evaluated statistical significance using a block-permutation null model. We extracted consecutive peak and valley lengths from the observed mask and independently shuffled them across permutations. During reconstruction of each permuted mask, we strictly maintained the alternating peak/valley order. Permuted profiles therefore followed either a peak–valley–peak–valley–... pattern or a valley–peak–valley–peak–... pattern; we did not permit adjacent regions of the same class, such as peak–valley–valley–peak. This generated randomized peak/valley profiles with the same block structure as the observed profile but with randomized spatial arrangement along the genome.

For each permuted track binary mask, we recalculated the enrichment statistic as:

$$S_{perm} = \sum_{i} (R_i M_i^{perm})$$

Where $R_i$ is the unchanged FGT ratio value at genomic position *i,* and $M_i^{perm}$ is the permuted peak/valley state of the tested track at that position. Repeating this procedure across 10,000 permutations generated a null distribution of enrichment statistics expected if the peak regions of the tested track were randomly distributed with respect to the FGT ratio profile.

We calculated empirical p-values as:

$$p = \frac{\#(S_{perm} \geq S_{obs}) + 1}{N + 1}$$

where N is the number of permutations, and $\#\left(S_{perm} \geq S_{obs}\right)$ is the number of permuted statistics greater than or equal to the observed statistic. The addition of one to both the numerator and denominator prevents zero p-values when no permuted statistic exceeds the observed statistic [6]. We considered associations significant when the empirical p-value was below 0.05, indicating that the FGT ratio signal accumulated within the peak regions of the tested track was greater than expected by chance under the block-permutation null model.

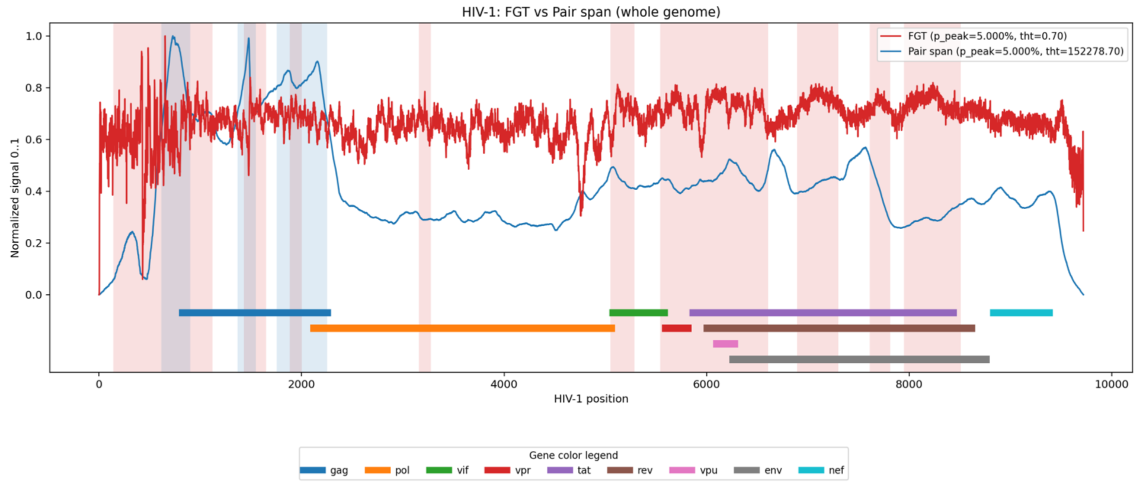


**Figure S1. HIV-1 FGT ratio profile compared with inter-polymorphisms interval coverage signal.** Genome-wide comparison between the HIV-1 FGT ratio profile and inter-polymorphisms interval coverage signal. The red line represents the FGT ratio profile, and the blue line represents the normalized inter-polymorphisms interval coverage track ("Pair span" in the figure), defined by cumulatively counting each genomic position whenever it falls between the two sites of an evaluated polymorphic-site pair. Shaded regions indicate peak intervals used in the permutation analysis, and HIV-1 gene annotations are shown. The block-permutation enrichment test did not detect a significant association between FGT ratio signal and inter-polymorphisms interval coverage regions (p=0.404).

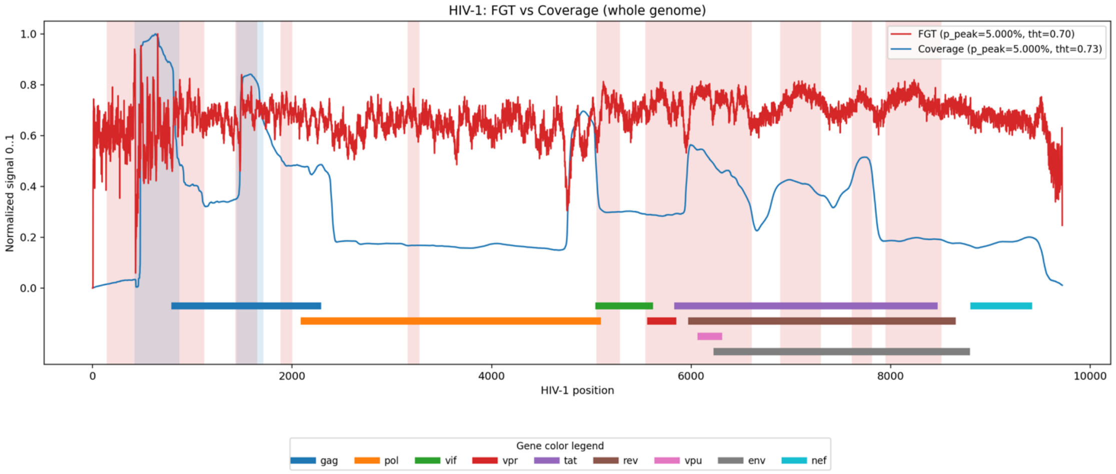


**Figure S2. HIV-1 FGT ratio profile compared with sequencing coverage.** Genome-wide comparison between the HIV-1 FGT ratio profile and sequencing coverage. The red line represents the FGT ratio profile, and the blue line represents the normalized coverage track, reflecting local read depth at each genomic position (only for reads that entered the analysis). Shaded regions indicate peak intervals used in the permutation analysis, and HIV-1 gene annotations are shown. No significant enrichment of FGT ratio signal was detected within high-coverage regions (p=0.788).

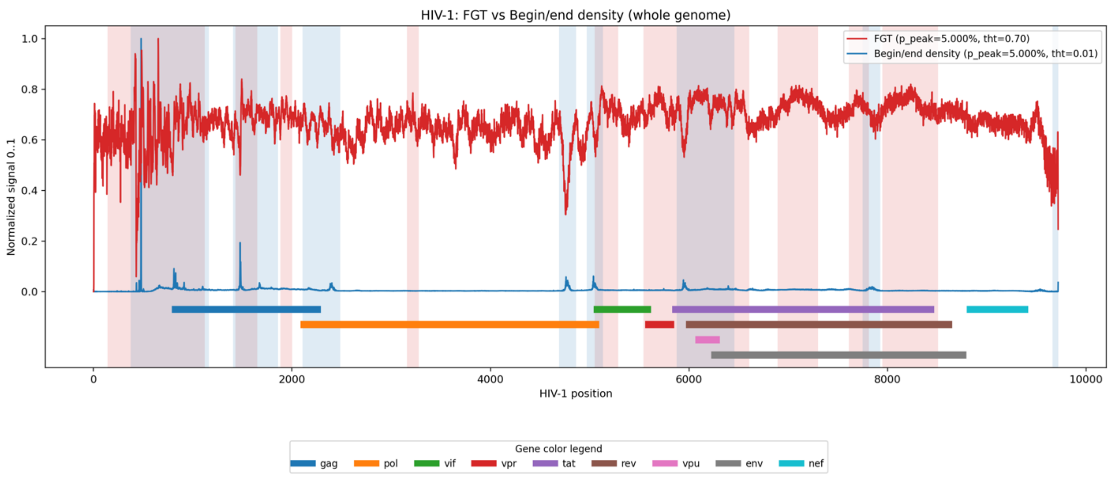


**Figure S3. HIV-1 FGT ratio profile compared with read start/end density.** Genome-wide comparison between the HIV-1 FGT ratio profile and read start/end density. The red line represents the FGT ratio profile, and the blue line represents the normalized read start/end density track, which reflects the accumulation of read starts and ends at each genomic position (only for reads that entered the analysis). Shaded regions indicate peak intervals used in the permutation analysis, and HIV-1 gene annotations are shown. The block-permutation enrichment test did not detect a significant association between FGT ratio signal and read start/end density peaks (p=0.595).

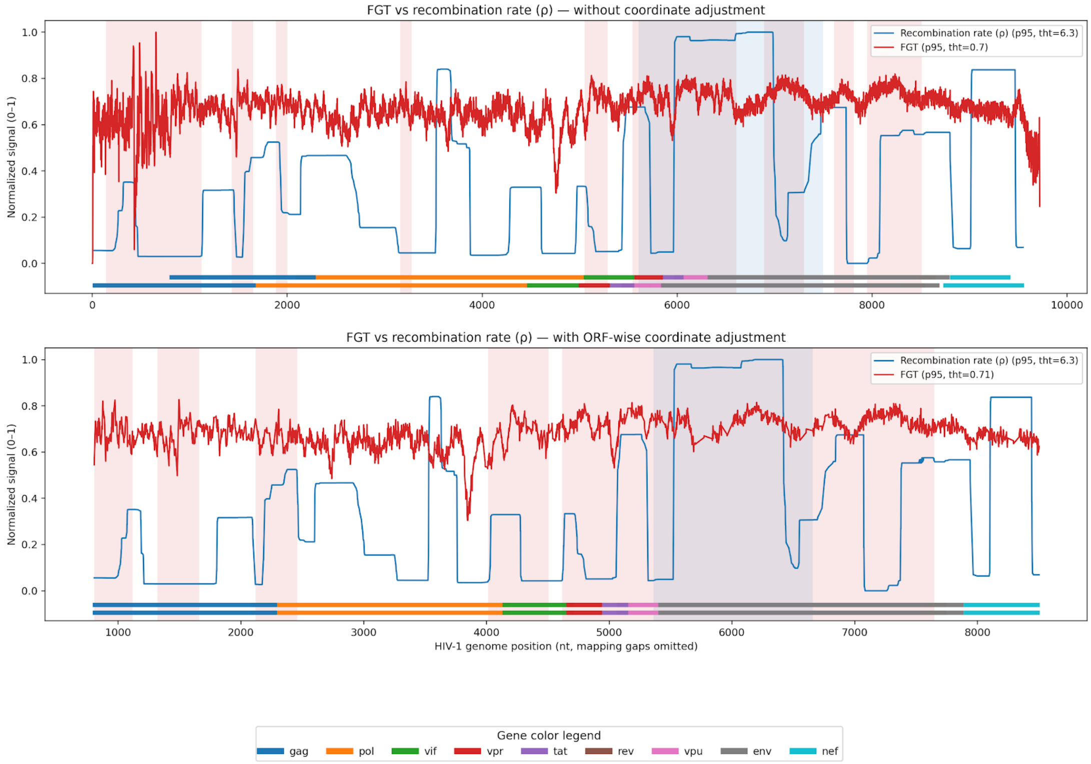


**Figure S4. HIV-1 FGT ratio profile compared with consensus sequence recombination rate estimates.** Comparison between the HIV-1 FGT ratio profile and the recombination rate (ρ) obtained with RDP5 from Simon-Loriere *et al.* [5] dataset. The red line represents the FGT ratio profile, and the blue line represents the recombination rate (ρ) along the genome. The upper panel shows the comparison without coordinate adjustment, whereas the lower panel shows the ORF-wise adjusted comparison. Shaded regions indicate peak intervals used in the permutation analysis, and HIV-1 gene annotations are shown. The block-permutation enrichment test detected a significant association between HIV-1 FGT ratio signal and the recombination rate obtained with RDP5 track after coordinate adjustment (p=0.046).

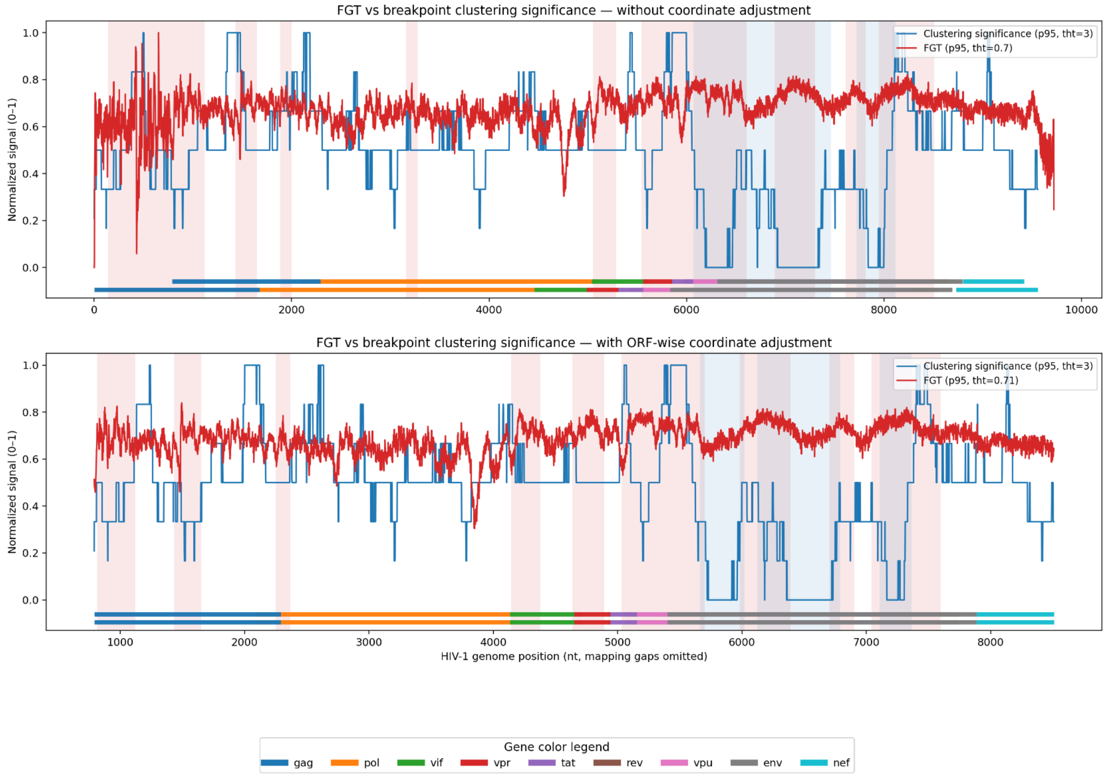


**Figure S5. HIV-1 FGT ratio profile compared with consensus sequence breakpoint-clustering significance.** Comparison between the HIV-1 FGT ratio profile and the breakpoint-clustering significance obtained with RDP5 from Simon-Loriere *et al.* [5] dataset. The red line represents the FGT ratio profile, and the blue line represents the breakpoint-clustering significance signal, indicating genomic regions where inferred recombination breakpoints showed stronger clustering. The upper panel shows the comparison without genomic coordinate adjustment, whereas the lower panel shows the ORF-wise adjusted comparison. Shaded regions indicate peak intervals used in the permutation analysis, and HIV-1 gene annotations are shown. The block-permutation enrichment test did not detect a significant association between HIV-1 FGT ratio signal and the breakpoint-clustering significance obtained with RDP5 track after coordinate adjustment (p=0.469).

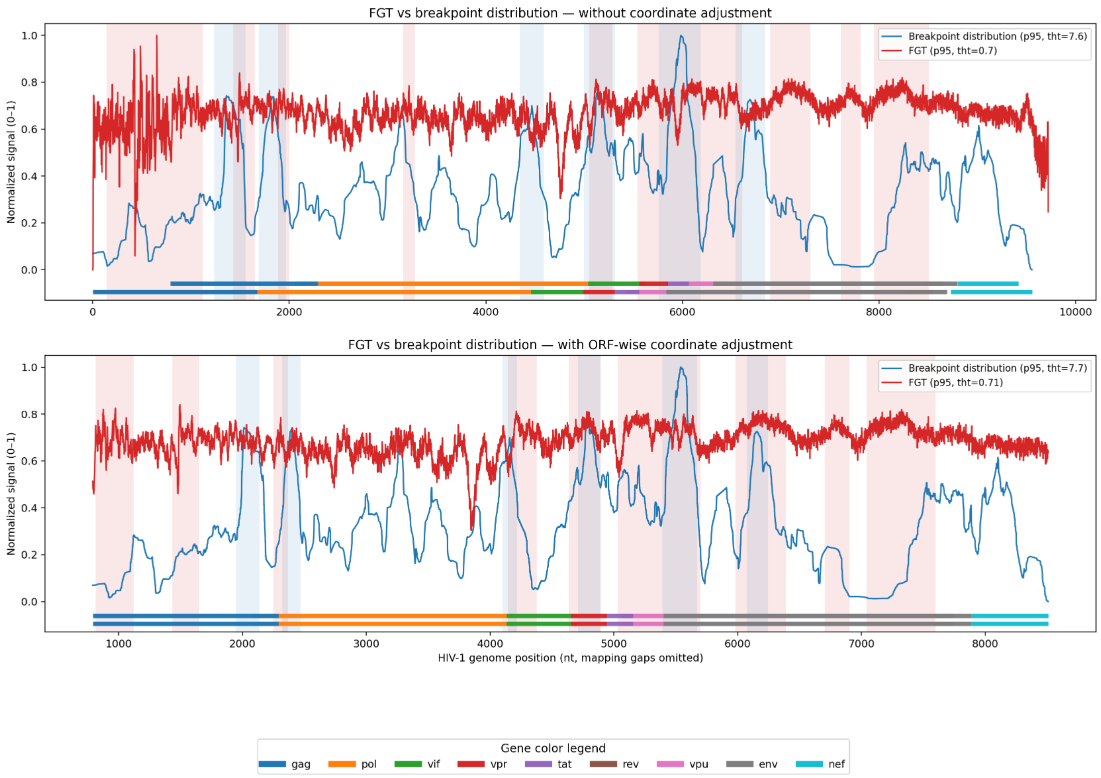


**Figure S6. HIV-1 FGT ratio profile compared with consensus sequence breakpoint distribution.** Comparison between the HIV-1 FGT ratio profile and the breakpoint-distribution track obtained with RDP5 from Simon-Loriere *et al.* [5] dataset. The red line represents the FGT ratio profile, and the blue line represents the breakpoint-distribution signal, indicating genomic regions where inferred recombination breakpoints accumulated more frequently. The upper panel shows the comparison without coordinate adjustment, whereas the lower panel shows the ORF-wise adjusted comparison. Shaded regions indicate peak intervals used in the permutation analysis, and HIV-1 gene annotations are shown. The block-permutation enrichment test detected a significant association between HIV-1 FGT ratio signal and the breakpoint-distribution obtained with RDP5 track after coordinate adjustment ($p = 0.005$).

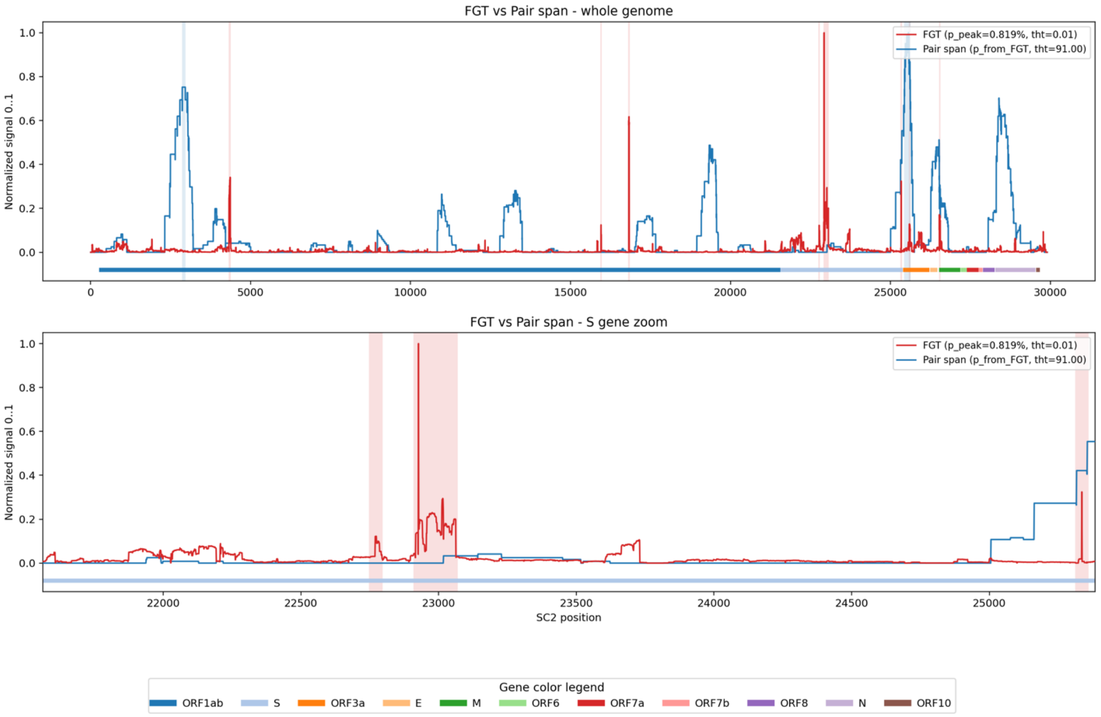


**Figure S7. SARS-CoV-2 FGT ratio profile compared with inter-polymorphisms interval coverage signal.** Comparison between SARS-CoV-2 FGT ratio profile and inter-polymorphisms interval coverage signal across the whole genome (upper panel) and the S gene (lower panel). The red line represents the FGT ratio profile, and the blue line represents the normalized inter-polymorphisms interval coverage track ("Pair span" in the figure), defined by cumulatively counting each genomic position whenever it falls between the two sites of an evaluated polymorphic-site pair. Shaded regions indicate peak intervals used in the permutation analysis, and gene annotations are shown. Although local overlap between FGT ratio and inter-polymorphisms interval coverage peaks was visible in the S-gene region, the block-permutation enrichment test did not support a significant genome-wide association (p=0.247).

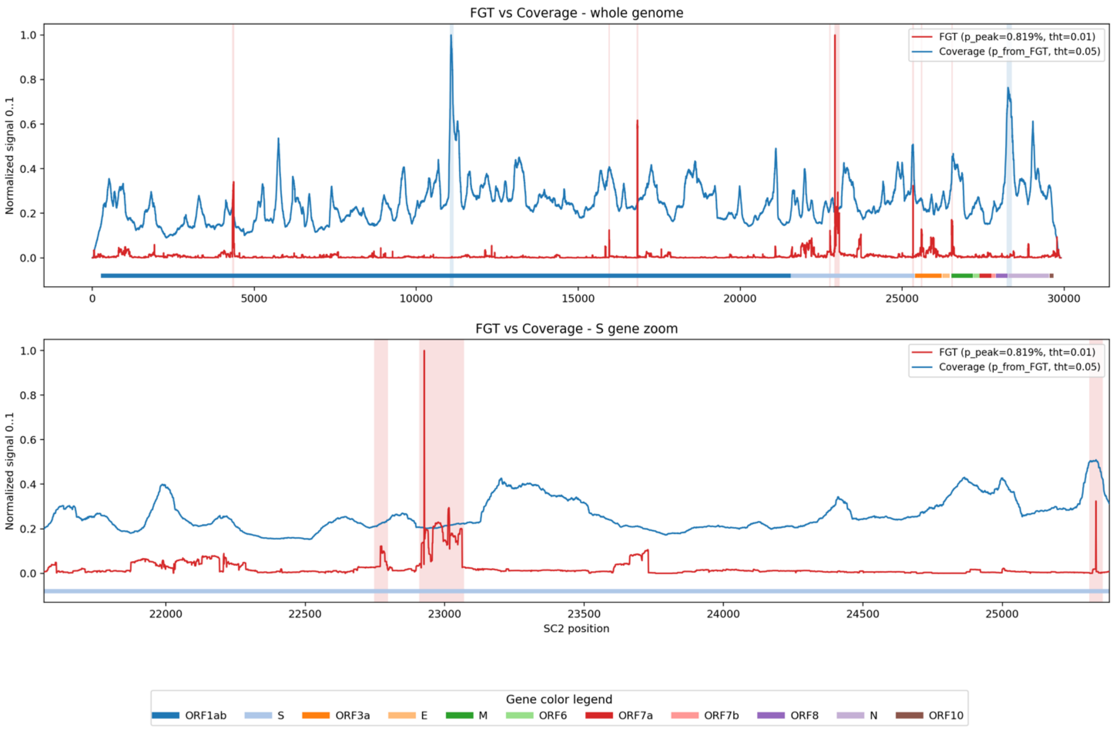


**Figure S8. SARS-CoV-2 FGT ratio profile compared with sequencing coverage.** Comparison between the SARS-CoV-2 FGT ratio profile and sequencing coverage across the whole genome (upper panel) and the S gene (lower panel). The red line represents the FGT ratio profile, and the blue line represents the normalized sequencing coverage track, reflecting local read depth at each genomic position (only for reads that entered the analysis). Shaded regions indicate peak intervals used in the permutation analysis, and gene annotations are shown. No significant enrichment of FGT ratio signal was detected within high-coverage regions (p=0.629).

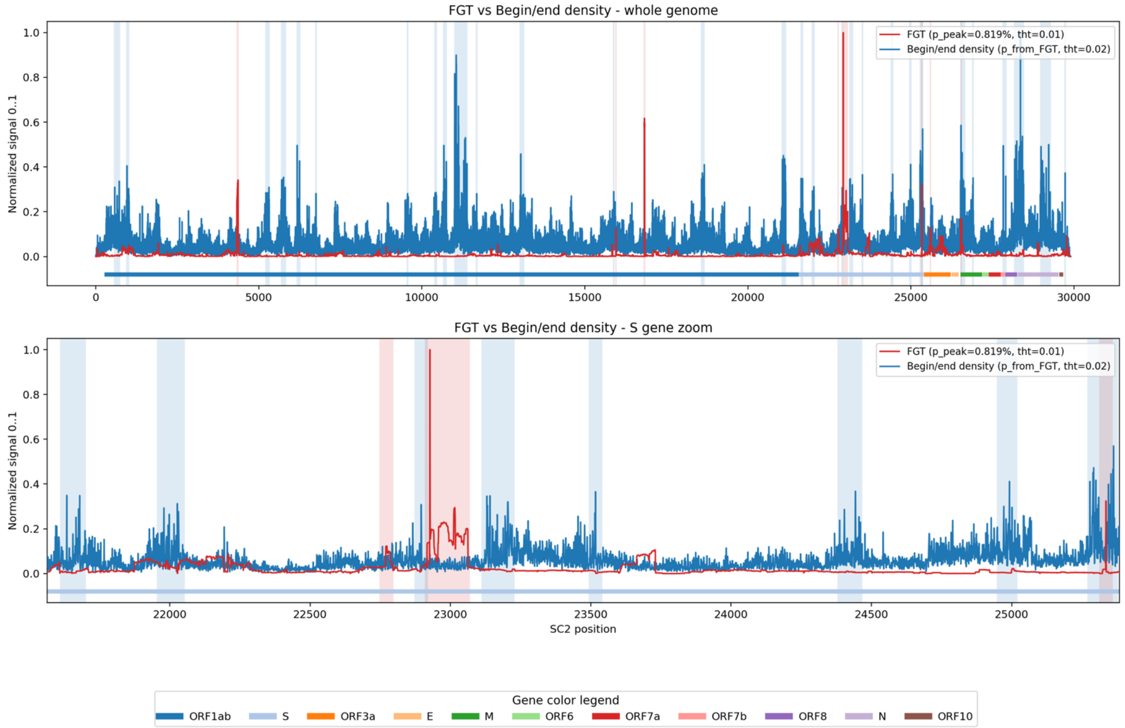


**Figure S9. SARS-CoV-2 FGT ratio profile compared with read start/end density.** Comparison between the SARS-CoV-2 FGT ratio profile and read start/end density across the whole genome (upper panel) and the S gene (lower panel). The red line represents the FGT ratio profile, and the blue line represents the normalized read start/end density track, which reflects the accumulation of read starts and ends at each genomic position (only for reads that entered the analysis). Shaded regions indicate peak intervals used in the permutation analysis, and gene annotations are shown. The block-permutation enrichment test did not detect a significant association between FGT ratio signal and read start/end density peaks (p=0.488).

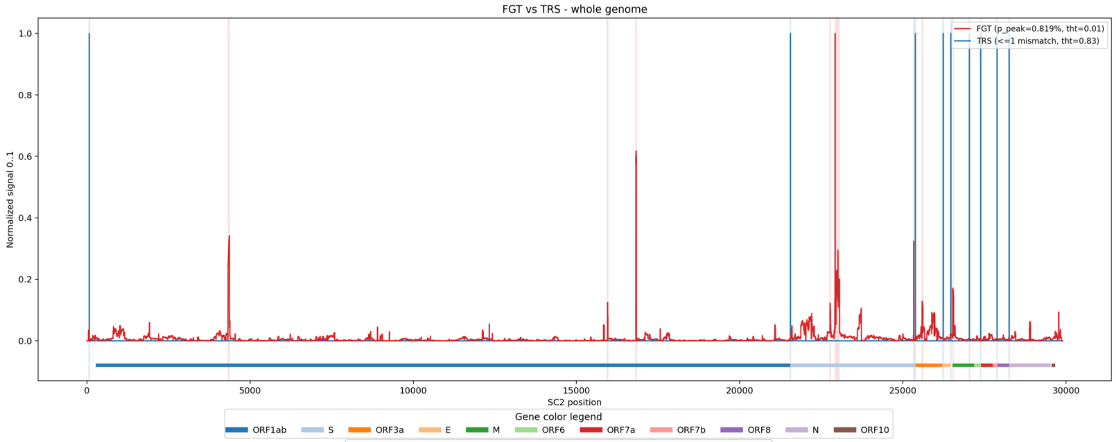


**Figure S10. SARS-CoV-2 FGT ratio profile compared with TRS-like positions.** Genome-wide comparison between the SARS-CoV-2 FGT ratio profile and TRS-like nucleotide positions. The red line represents the FGT ratio profile, and the blue line indicates TRS-like positions, defined by allowing up to one mismatch from the TRS motif (ACGAAC). Shaded regions indicate peak intervals used in the permutation analysis. Gene annotations are shown. The block-permutation enrichment test did not detect a significant association between FGT ratio signal and TRS-like positions ($p=0.597$).

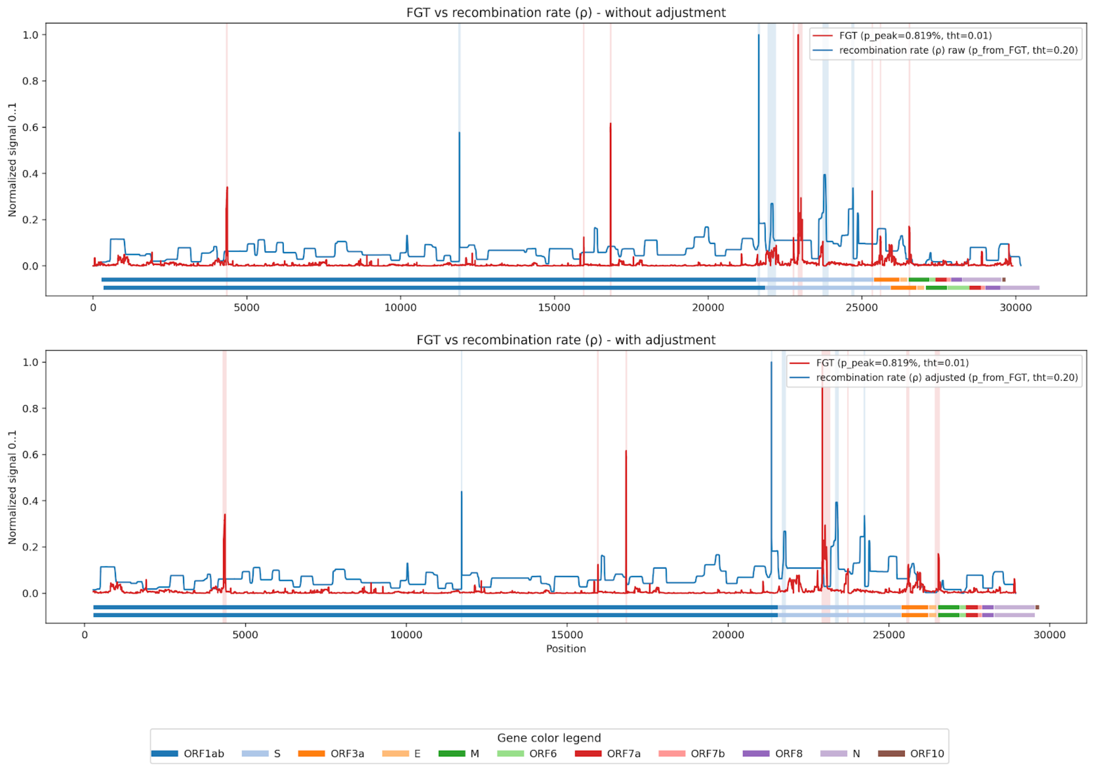


**Figure S11. SARS-CoV-2 FGT ratio profile compared with Sarbecovirus recombination rate (ρ).** Comparison between the SARS-CoV-2 FGT ratio profile and the Sarbecovirus-derived recombination rate (ρ) track extracted from the RDP5 analysis from De Klerk *et al.* [4] dataset. The red line represents the FGT ratio profile, and the blue line represents the Sarbecovirus recombination rate (ρ), indicating genomic regions with higher inferred recombination signals across Sarbecovirus. The upper panel shows the comparison without genomic coordinate adjustment, whereas the lower panel shows the adjusted comparison. Shaded regions indicate peak intervals used in the permutation analysis, and gene annotations are shown. The block-permutation enrichment test did not detect a significant association between SARS-CoV-2 FGT ratio signal and the Sarbecovirus recombination rate track (p=0.405).

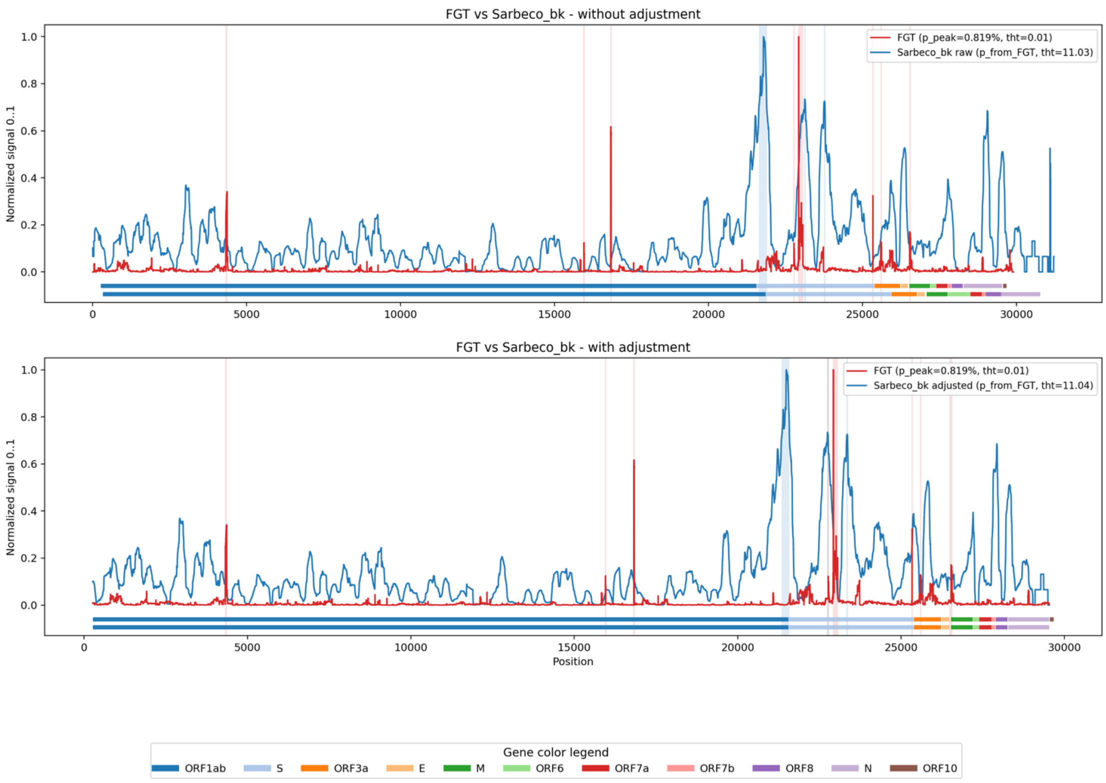


**Figure S12. SARS-CoV-2 FGT ratio profile compared with Sarbecovirus breakpoint-density signal.** Comparison between the SARS-CoV-2 FGT ratio profile and the Sarbecovirus-derived breakpoint-density track extracted from the RDP5 analysis from De Klerk *et al.* [4] dataset. The red line represents the FGT ratio profile, and the blue line represents the Sarbecovirus breakpoint-density signal, indicating genomic regions where recombination breakpoints were more frequently inferred across Sarbecovirus. The upper panel shows the comparison without genomic coordinate adjustment, whereas the lower panel shows the adjusted comparison. Shaded regions indicate peak intervals used in the permutation analysis, and gene annotations are shown below the genome coordinate axis. No significant enrichment of FGT ratio signal was detected within Sarbecovirus breakpoint-density peak regions (p=0.548).
